# Crossing effects in the tactile temporal order judgment task: A meta-analysis

**DOI:** 10.1101/2025.05.05.652194

**Authors:** Nina Schmelter, Benedikt Langenberg, Axel Mayer, Tobias Heed

**Author notes:** Correspondence T.H. Author Contributions Contributions according to https://credit.niso.org NS: conceptualization, data curation, formal analysis, investigation, methodology, validation, visualization, writing – original draft BL: formal analysis AM: conceptualization, methodology, formal analysis, project administration, supervision, validation, writing – original draft TH: conceptualization, project administration, supervision, writing – original draft.

## Abstract

The tactile temporal order judgment (TOJ) task is widely used in multisensory neuroscience. Participants judge which of two tactile stimuli, one on each hand, came first. A key finding is that TOJ performance declines when the arms are crossed, likely due to interactions between tactile, proprioceptive, and visual information. The TOJ crossing effect has been widely reported. However, studies have used different analysis methods, leaving open whether the choice of method influences the effect’s magnitude. Some studies have also reported that the crossing effect is modulated by moderators such as the analysis method, availability of visual information, response modality, or speeded response requirements. Through a systematic, exhaustive literature search, we identified 85 experiments in 51 publications that investigated the TOJ crossing effect, of which 39 experiments from 29 publications reported sufficient information for inclusion in a meta-analysis and fit the focus of our study. This analysis estimated the crossing effect’s size at approximately 1.45. Checks for potential publication bias indicated that this estimate may be inflated, with a corrected estimate of 0.97 using a regression-based adjustment for small-study effects (Rücker’s method), and 1.4 based on p-curve analysis of the reported p-values. Although analysis method and response mode were identified as significant moderators, most moderators and specific contrasts were not statistically significant, likely because too few studies were available to achieve adequate statistical power. The large effect size supports the TOJ task’s use in other research, e.g. developmental and clinical. However, the lack of significant moderation suggests caution when applying the task to questions beyond the crossing effect itself.

**Public Significance Statement:** We provide an effect size estimate of a well-known tactile-spatial measure, the tactile temporal order judgment crossing effect, based on a PRISMA-guided literature review. This measure will help planning for sufficient power in future studies.

## Introduction

Crossing the arms over the body midline impairs tactile choice tasks, a phenomenon known as the tactile crossing effect [1]. A prominent paradigm for its investigation is the tactile Temporal Order Judgment (TOJ) task, in which participants report which of two consecutive stimuli, one applied to each hand, occurred first (or second), typically by pressing a button with the respective hand [2,3]. Intuitively, arm posture should be irrelevant to choosing which hand to press a button. However, when participants cross their arms, TOJ performance is strongly impaired and can even be systematically reversed. As a result, the TOJ task has become a prime example of a tactile crossing effect and has been used extensively over the last two decades to investigate tactile spatial processing.

The origin of crossing effects such as that observed in the TOJ task has been debated. The most prominent idea is that they arise from a conflict between different spatial codes: a skin-based, or anatomical, code that signals where on the skin the touch occurred, and a spatiotopic, or external, code that signals where the touch was in space [1,4]. With crossed arms, these two codes carry opposing information; for instance, the right hand is then located in left space, and this kind of conflict may impair tactile judgments. Newer studies have cast doubt on this account and suggest that crossing effects instead stem from confusion of the limb’s default posture –where it is usually located– and its actual position at the moment of touch [5–7].

Irrespective of its theoretical underpinnings, the TOJ crossing effect is often large, and this likely contributes to the task’s popularity. However, there are multiple versions of the TOJ paradigm, and the task has been analyzed in multiple ways. This variability in implementation and analysis makes it difficult to compare studies directly, beyond qualitative similarities. Moreover, no prior work has estimated a statistical effect size based on evidence from multiple studies.

An effect size estimate would aid future research, for example, in planning sample sizes based on statistical power or in defining priors in Bayesian analysis. The TOJ task has been employed in developmental and clinical studies, including research on mental disorders, physical illnesses, and brain processes [8–11]. Such studies typically aim to optimize experimental efficiency and determine the minimum number of required participants. One option to reduce experimental time is to use an adaptive version of the TOJ task, that is, online, trial-by-trial adjustment of testing parameters to expedite reliable performance estimation [12]. Still, a proper estimate of the expected effect size is necessary to determine an appropriate sample size. As a result, experiments are often either inefficient or underpowered [13].

The TOJ task has been employed for a wide range of scientific questions, and numerous manipulations have been applied next to limb crossing. Some of these have been examined across multiple studies, potentially allowing estimating their effect sizes as well. One key variable that has differed across studies is the response modality. Often, participants have responded with the hands, that is, with the same limbs that received stimulation. Some studies, however, required button responses with the feet. This response modality requires instructions for mapping the stimulated hand to the responding foot: under anatomical instructions, responses are given with the foot on the same body side as the hand that received the first stimulus; In contrast, under external instructions, responses are given with the foot on the same side of external space as the stimulated hand. These two instructions require different responses when the hands are crossed: if the right hand was first, the participant presses the right foot under anatomical instructions (stimulus on the right body side) but the left foot under external instructions (stimulus in left external space due to the crossed posture). Several studies have reported a larger TOJ crossing effect for external than anatomical instructions [14,15]. In yet other studies, participants responded verbally [11,16], but, to our knowledge, no direct comparison of verbal vs. button-press responses has been published.

A second manipulation investigated in multiple studies is the availability of visual information. The crossing effect was reported to be diminished when the hands were crossed behind the back [17], when the eyes were closed [18], and when uncrossed rubber arms were placed above the hidden, crossed arms [19]. Although most studies have reported that the absence of visual input reduces the crossing effect, some findings have been inconsistent [20].

A third aspect which may affect TOJ is task instructions regarding speed and accuracy. The TOJ has sometimes been run as an unspeeded task that emphasizes correct responding [2], whereas many studies have asked participants to respond both as fast and accurately as possible [3,21–23]. Emphasizing speed may compel participants to respond before they have fully processed the stimuli, leading to more frequent guessing and errors in the crossed posture, which may, in turn, amplify the crossing effect. We are not aware of any study that systematically addressed this issue for the TOJ task.

Finally, the reported size of experimental effects may systematically depend on the analysis method [1]. TOJ performance is often assessed using a range of stimulus onset asynchronies (SOAs) between the two tactile stimuli. By convention, SOAs are marked as negative when the left hand stimulation occurred first and plotted as percent “right-first” responses (y-axis) against the SOA (x-axis; see Fig. 1 for examples). With uncrossed hands, performance follows a typical psychophysical, S-shaped curve. In contrast, with crossed hands, the curve is often N-shaped: responses for long SOAs are correct, but responses for short SOAs are reversed [ref 3, see Fig. 1A]. This N-shaped result pattern is unusual; to our knowledge, it is not observed in other psychophysical tasks. Reaction time (RT) increases as the SOA decreases and is significantly longer in the crossed than in the uncrossed posture [3,23].

**Figure 1.**
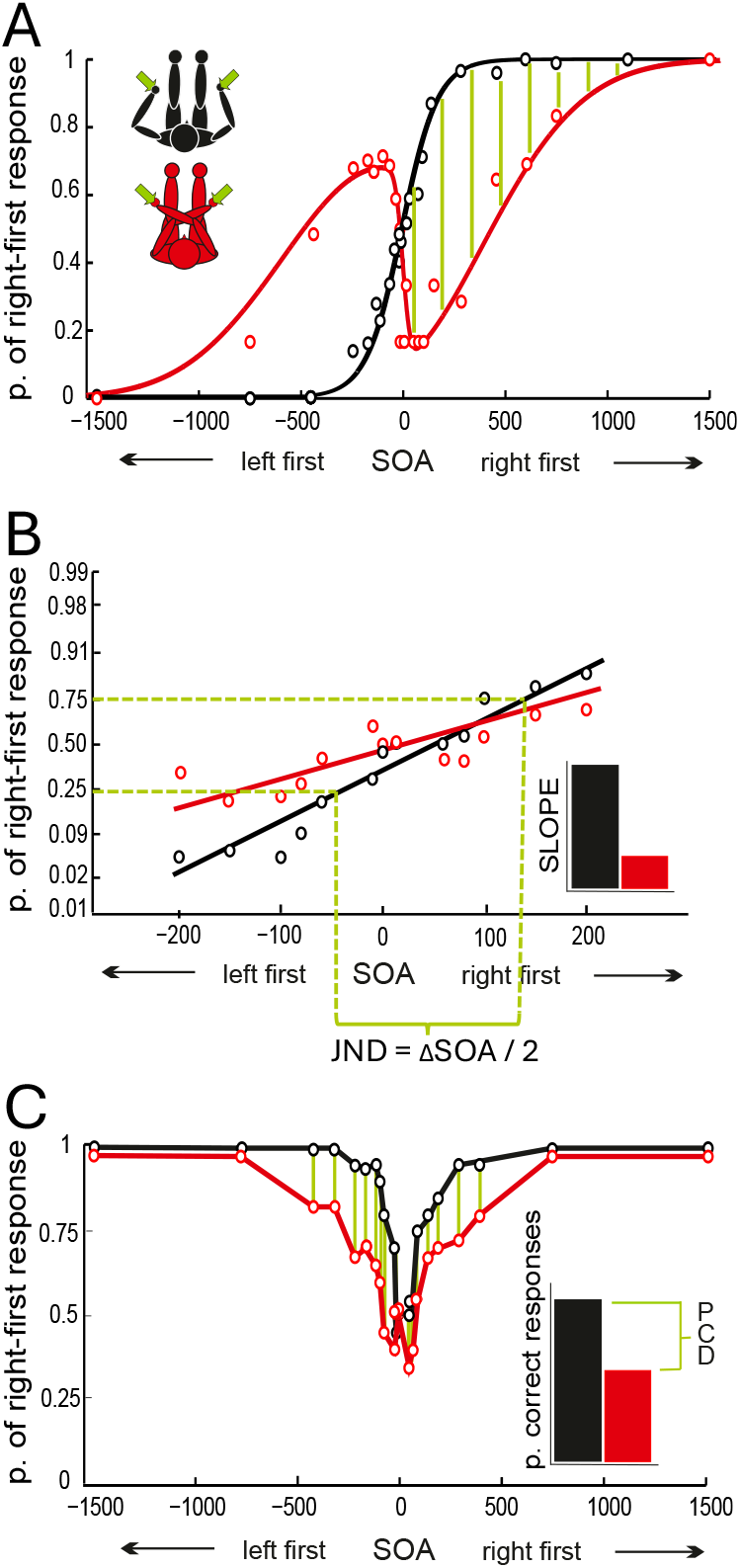
Analysis variants for TOJ crossing effects. In most TOJ experiments, the two tactile stimuli are presented with multiple stimulus onset asynchronies (SOAs), and a global performance measure is derived across SOAs. Black traces in all panels reflect performance with uncrossed hands; red traces reflect performance with crossed hands. (A) Illustration of the systematically reversed responses often observed for tactile TOJ with crossed hands. The flip model analysis fits the difference between uncrossed and crossed performance, quantified as the probability of right-first responses, for left-first and right-first stimulus pairs with Gaussians. This difference is illustrated here for right-first responses with green hatching; see [3] for illustration of the fit. (B) Regression of probit-transformed right-first response probability. Both the regression line’s slope and half the difference between the x-values corresponding to 25% and 75% correct responses have been used as performance measure [2]. (C) Illustration of the proportion-correct difference, which sums the accuracy difference between the uncrossed and crossed conditions, illustrated by the green lines, across all SOAs [21]. The figure has been modified from [1].

Whether speeded or not, most reports have focused on analyzing TOJ accuracy rather than RT. One approach has been to probit-transform the percentage of right-first responses to linearize the S-shaped response curve and to derive the slope of the resulting straight line (ref. [2], see Fig. 1B). A second measure has been the just noticeable difference, defined as half the distance between the SOA required for 25% and 75% correct right-first responses after probit transformation (ref [2], see Fig. 1B). A third measure has been to sum accuracy across all SOAs (ref. [21], see Fig. 1C). With all three methods, the crossing effect was defined as the difference of the respective measure between the uncrossed and crossed postures. A fourth analysis method has been to fit the N-shaped response curves observed in the crossed posture (see Fig. 1A). This can be done by integrating three Gaussian components: the cumulative Gaussian fitted to uncrossed performance (see Fig. 1A, black trace) and two regular Gaussians that fit the difference between uncrossed and crossed performance for left-first and right-first stimulus pairs (illustrated for right-first stimulus pairs in Fig. 1A; see ref. [3] for details). The two latter two Gaussians fit the lobes of the N, which reflect flipped–that is, systematically reversed and, thus, incorrect–responses in the crossed posture. Their amplitudes, termed A_right_ and A_left_, are then interpreted as a measure of the crossing effect [3,24,25]. The two “flip Gaussians” are derived solely for the crossed condition; this distinguishes the flip model’s fitting procedure from all other crossing effect measures, which all rely on subtracting identically computed performance measures between uncrossed and crossed conditions. Nonetheless, the crossed posture’s flip model fit is based on the individual’s uncrossed posture’s fit, so that the flip Gaussians capture the difference between the two postures. The flip model also allows deriving JNDs, but 16%/84% correct right-first have been used in this case, owing to the model fitting a Gaussian for the uncrossed condition, and using a one standard deviation cut-off yielding these percentages [3,26].

There is no consensus on which analysis method is most appropriate, and no method has been shown to be generally more sensitive or to consistently yield a larger crossing effect [23]. In addition to these analysis choices, the selection of the SOAs used in the experiment, as well as some further analysis options, may result in differences across studies [27].

In sum, both the need for an effect size estimate to aid future study planning, the lack of consensus about the formal analysis of the TOJ crossing effect, and the unsystematic exploration of potential moderators, all call for a systematic review. We address these issues by conducting a meta-analysis for the TOJ crossing effect and its moderation by response mode, visual input, speed and accuracy instructions, and analysis method.

## Methods

### Ethics, Data availability statement, Preregistration, Transparency, and Openness

No ethical approval was sought for the present study because it is based entirely on summarizing existing data. Below, we report how we determined our sample size (i.e., selected the papers for our meta-analysis), all data exclusions, all manipulations, and all measures. All variables created for our report, as well as the R code used for analysis, are freely available at https://osf.io/xtd9g/. The data set also includes several variables extracted but not used in our report, such as the reported *p* and *df* values for *t, F*, and ***χ***^2^ tests. We did not preregister this study. Our report adheres to the TOP Research Practices (see https://www.cos.io/initiatives/top-guidelines), with Level 1 (Disclosure) for Study Registration and Level 2 (Shared and Cited) otherwise.

### Search Strategy

We searched the databases BASE, EMBASE, Pubmed, and Web of Science, accessed through Bielefeld University on April 28, 2023. We chose these databases because they have demonstrated high levels of precision, recall, and reproducibility [28]. We employed the search string “(tactile OR hand OR arm) temporal order judg*” for all databases, using the appropriate syntax for each (see Appendix A, Table A1). We included only English articles when this option was available. The search and article screening were performed by NS and discussed with TH and AM.

### Selection Criteria

We included studies in the meta-analysis if they: (1) conducted the tactile TOJ task with crossed and an uncrossed conditions, (2) presented stimuli sequentially to both hands, (3) asked participants to judge the temporal order of the two stimuli, (4) collected responses via the hands, feet, or verbally, (5) included participants aged 18 and older, (6) were published peer-reviewed publications, (7) were written in English.

We excluded studies if they: (1) reported mental or health impairment of participants and did not include an unaffected control group; or (2) did not provide sufficient data within the published article to calculate a crossing effect.

### Data Extraction and Derived Variables

We collected all studies in Zotero (Corporation for Digital Scholarship, Vienna, Virginia, USA, https://www.zotero.org). We used automated procedures to eliminate duplicates and removed any remaining duplicates manually. One coder (NS) assessed the remaining studies for inclusion and extracted data from each study, including statistical information to calculate effect sizes and standard errors. The same coder double-checked all results to ensure accuracy.

Additionally, we operationalized potential moderator variables and coded them for each study as follows:

1. Analysis method for accuracy
  a. “accuracy” – the difference in performance between the crossed and uncrossed condition was calculated for each SOA and then pooled over all SOAs, as introduced in [21]
  b. “slope” – the S-shaped curve of the percentage of right-first responses was probit, logit, or Weibull transformed to linearize the psychometric curve, and the resulting straight line displayed the performance in the TOJ task as a function of the SOA
  c. “JND 75%” – the JND was obtained from linearized response data (see b) and calculated as the time window between the two applied stimuli in which 75% of the judgments were correct)
  d. “JND 84%” – response data were pooled across participants and fitted with a cumulative four-parametric cumulative Gaussian function, as introduced in [3]. Subsequently, the standard deviation (σ) was calculated, reflecting the smallest time window between the two applied stimuli in which 84% of the judgments were correct)
  e. The fitted parameter A from the flip model [3], averaged across the two estimates A_left_ and A_right_ If multiple outcome measures were available, we entered only one into our analysis, using the following order: accuracy, slope, JND 75%, JND 84%, flip model’s A.
2. visual input
  a. “eyes open” – participants’ eyes were open. We did not differentiate whether fixation was required or not; it is known that gaze direction influences TOJ; however, the reported effect was very small compared to the general crossing effect [4]. Moreover, we did not differentiate whether participants’ hands were hidden or not; we are not aware of any study having tested this differentiation previously. Moreover, we have not encountered this differentiation as a scientific question in papers on tactile TOJ or at conferences and therefore opted to ignore it here.
  b. “eyes closed” – participants were blindfolded
3. response modality and mapping
  a. “hands” participants responded with the hand or a finger of the hand that was stimulated first [or second]
  b. “feet external” – participants responded with the foot on the same side of space as the hand stimulated first [or second]
  c. “feet anatomical” – participants responded with the foot on the same body side as the hand stimulated first [or second]
  d. “verbal” participants responded verbally
4. speed/accuracy instructions
  a. “unspeeded” – participants responded without any time restriction and/or were instructed to prioritize accuracy
  b. “time-restricted” – participants had to respond within a specific time interval or were instructed to respond as accurately and as quickly as possible; the latter were usually combined)

We did not conduct a formal risk of bias assessment, reasoning that the TOJ task is typically performed in controlled laboratory settings.

### Statistical Analysis

#### Effect Measures

All included studies used within-subject designs, and all but two aggregated the dichotomous dependent variable (correct or incorrect) into a percentage correct score that can be treated as continuous for the range of values that is not too close to the limits. Therefore, we calculated a version of Cohen’s *d* known as Cohen’s *d*_*z*_ or *d*_*D*_ (hereafter referred to as Cohen’s *d*_*z*_), which is appropriate for within subject-designs with a continuous outcome. Using Cohen’s d_z_ allowed us to pool primary studies with different outcome measures, such as slope vs. percentage correct. One possibility to calculate Cohen’s *d*_*z*_ is to compute the mean difference between crossed and uncrossed conditions and then use the standard deviation of these difference scores as the denominator [29]:

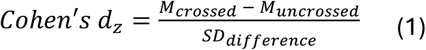

The mean difference and its standard deviation were only available in studies that used accuracy as an outcome measure. Alternative derivations are available for Cohen’s *d*_*z*_ depending on the available information:

a. calculation based on the *t*-value and the sample size [30]:

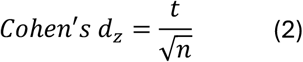
b. calculation based on the *F*-value (with one numerator degree of freedom) and the sample size [31]:

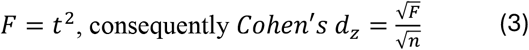
c. from the *χ*^2^-value and the sample size, with the *χ*^2^-value for categorical outcomes treated as *F*-value to obtain comparability with the analysis of continuous variables as described in b:

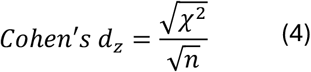
d. from the regression coefficient (β) in a generalized linear mixed model (GLMM) and the sample size, employing the additonal assupmtions that the β-value is unstandardized, and the *z*-value is approximately equivalent to the *t*-value:

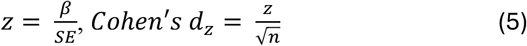
e. from the flip model’s fitted parameter A, averaged across A_left_ and A_right_, by regular normalization:

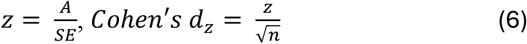 We assessed whether this strategy is reasonable by calculating d_z_ according to equation (6) for two experiments published by the last author (Exp 1, condition CONTROL, of ref. [23]; Exp. 2, condition “All Straight”, of ref. [4]). The effect sizes were 1.00 and 1.27, respectively. As these values fell within the range of estimates obtained with other measures, we proceeded using equation (6) for our meta-analysis. We did not include these two test values into our meta-analysis because we did not compute any effect sizes from raw data for any other measure either.
f. from the means of the crossed and uncrossed conditions, their standard deviations, and the correlation between the two conditions [29]; we estimated the required correlation from three openly available datasets (see Supplementary Information for details):

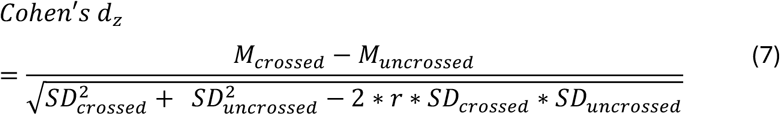

Equations (4) and (5) are approximations based on the properties of *F* and *t*, which converge against the *χ*^2^ and the normal distribution for large *n*.

If multiple statistical metrics were available to calculate the effect size, formulas were chosen in the order reported above.

#### Standard Error

We calculated the standard error of each effect size according to equation (7); see Appendix B for its derivation.

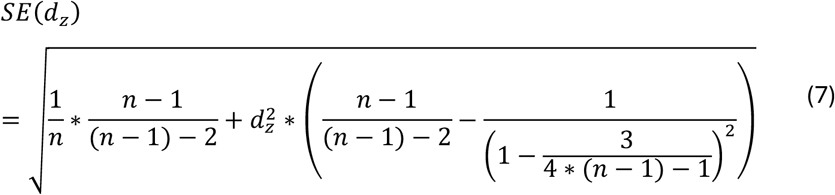

#### Meta-Analysis

We conducted statistical analyses with R [32] and R Studio [33]. We employed the packages meta [34] to conduct the meta-analysis, dmetar [35] for outlier and influence analysis, metasens [36] for Rücker’s Limit Meta-Analysis Method, tidyverse [37] for data handling and multcomp [38] for post-hoc pairwise comparisons.

We used a random-effects model to pool effect sizes, because we anticipated considerable between-study heterogeneity. We calculated the heterogeneity variance (*τ*^2^) using the restricted maximum likelihood (REML) estimator [39] and the confidence interval around the pooled effect using Knapp-Hartung adjustments [40].

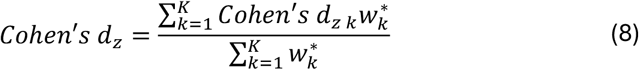

where:

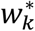 is the weight assigned to each study *k*

K is the number of studies included in the meta-analysis

with:

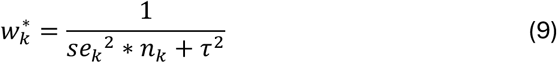

where:

*n*_*k*_ is the sample size of each study *k*

We used Cochran’s Q to test if between-study variation exceeded sampling error and estimated the proportion of variability due to true differences using Higgins and Thompson’s *I*^2^ [41]. Additionally, we calculated prediction intervals to estimate the expected range of effects in future studies [42]. To detect divergent studies, that is, studies that differ markedly from the study pool, we set as outlier criterion that a study’s 95% confidence interval was beyond the 95% confidence interval of the pooled effect. Additionally, we identified studies with a high influence on the pooled effect size using the leave-one-out method. To this end, we calculated the difference for Cohen’s *d*_*z*_ with and without each individual experiment. The resulting differences were plotted in a Baujat plot on the y-axis, with overall heterogeneity, measured by Cochran’s Q, on the x-axis [43].These differences were plotted as a Baujat plot, in which Cochran’s Q (i.e., the experiment’s contribution to overall heterogeneity) is plotted on the x-axis, the experiment’s influence on the pooled effect (i.e., the absolute change in the effect size estimate from the leave-one-out analysis in units of Cohen’s *d*_*z*_) is plotted on the y-axis.

#### Analysis of moderator effects

To explore the impact of the four potential moderators of the crossing effect’s effect size–analysis method, visual input, response mode, and speed and accuracy instructions– we conducted separate moderator analyses, each restricted to studies that reported the relevant moderator. Ten studies lacked information on one moderator each; therefore, we used separate analyses to retained as many studies as possible for each moderator [44]. We fit random-effects (plural) models for these analyses [45]. When any subset contained fewer than six studies, we used a pooled estimate of the between-study heterogeneity variance, *τ*^2^, computed across all studies subsets [46]. When a moderator was significant, we conducted post-hoc pairwise comparisons by fitting a meta-regression model without an intercept term and then computing Tukey’s post-hoc tests for contrasts of all subgroup pairings.

#### Assessment of potential publication bias

We employed small-study effect methods to assess potential publication bias. These methods assume that studies with smaller sample sizes are more susceptible to bias. As sample size decreases, standard errors increase, inflating the variability of estimated effect sizes. Consequently, only comparatively large effects are likely to reach statistical significance in small studies. If publication is more likely when results are significant, smaller studies that, by chance, yield large effect sizes (and thus significant *p*-values) become overrepresented. Small-study effect methods evaluate whether the published literature shows this pattern by testing whether smaller studies report disproportionately larger effects [47].

We assessed small-study effects in three steps. First, we visually inspected a funnel plot [34,48,49]; in the absence of publication bias. Effect sizes should be symmetrically distributed around the meta-analytic mean. In a second step, we computed an adjusted effect size with Rücker’s limit meta-analysis method [36,50]. Finally, in step three, we applied p-curve tests for right-skewness and flatness [47]. Unlike methods targeting small-study effects, a p-curve analysis examines the distribution of reported p-values across studies. In the absence of a true effect, p-values should be uniformly distributed, yielding a flat curve. Selective reporting and practices such as p-hacking, however, can produce a left-skewed distribution, that is, a higher-than-expected proportion of p-values just below the significance level. By contrast, a true effect without publication bias leads to a right-skewed p-curve, with smaller p-values occurring more frequently. Skewness increases with effect size and sample size. Given the large effects in the present analysis, pronounced right-skewness would be expected in the absence of bias. We used the right-skewness test to evaluate the null hypothesis that no effect exists (i.e., a flat distribution), and the flatness test to evaluate whether a small effect exists.

## Results

### Search Results

The database search yielded 973 publications (Fig. 1), which reduced to 346 eligible publications after duplicate removal. We excluded 301 publications based on abstract screening and fully reviewed 45 publications comprising 85 experiments. Of these, we excluded 16 publications (49 experiments): reported data were insufficient for meta-analytic inclusion for 16 experiments, and at least one inclusion criterion was not met by the other 33 experiments. We list the references to the fully evaluated publications and the reasons for their exclusion in Supplementary Table B1.

The remaining 29 publications comprised 39 experiments with a total of 688 participants (see Table 1 for references and study details). We entered data as separate experiments when a publication reported multiple experiments with distinct participant samples, and when a between-subject factor created independent groups (e.g. some participants tested with eyes open, some with eyes closed).

**Table 1.** Included studies and characteristics used for the meta-analysis.

| **Author** | ***n*** | **Statistical characteristics** | **Outcome measure** | **Eyes** | **Response modality** | **Speed** |
| --- | --- | --- | --- | --- | --- | --- |
| Azañón & Soto-Faraco (2007) [19] - exp. 1 | 19 | *M, SE* | JND 84% | open | hands | unspeeded |
| Azañón & Soto-Faraco (2007) [19] - exp. 2 | 20 | *M, SE* | JND 84% | open | hands | unspeeded |
| Azañón et al. (2015) [52] | 12 | *F* | JND 75% | open | hands | unspeeded |
| Azañón, Mihaljevic et al. (2016) [53] | 16 | *t* | JND 75% | closed | hands | unspeeded |
| Azañón, Radulova et al. (2016) [54] | 11 | *t* | slope | NA | hands | unspeeded |
| Badde et al. (2014) [55] - exp. 1 | 17 | *F* | Accuracy | open | hands | time restriction |
| Badde et al. (2014) [55] – exp. 2 | 17 | *F* | Accuracy | open | hands | time restriction |
| Badde et al. (2015) [56] - exp. 1 | 13 | *F* | Accuracy | open | hands | unspeeded |
| Badde et al. (2015) [56] - exp. 2 | 14 | *F* | Accuracy | open | hands | unspeeded |
| Badde et al. (2017) [57] | 16 | $\chi^2$ | Accuracy | NA | hands | time restriction |
| Cadieux & Shore (2013) [14] - eyes closed, feet external | 11 | *t* | Accuracy | closed | feet external | time restriction |
| Cadieux & Shore (2013) [14] - eyes closed, hands | 20 | *t* | Accuracy | closed | hands | time restriction |
| Cadieux & Shore (2013) [14] - eyes open, feet external | 11 | *t* | Accuracy | open | feet external | time restriction |
| Cadieux & Shore (2013) [14] - eyes open, hands | 20 | *t* | Accuracy | open | hands | time restriction |
| Cadieux et al. (2010) [21] | 48 | *t* | slope | open | feet external | time restriction |
| Crollen, Albouy, et al. (2017) [58] | 11 | *t* | slope | closed | hands | time restriction |
| Crollen, Lazzouni, et al. (2017) [22] | 11 | *t* | slope | closed | hands | time restriction |
| Crollen et al. (2019) [59] | 15 | *t* | slope | closed | feet external | time restriction |
| Heed et al. (2012) [23] - exp. 1 | 15 | *F* | Accuracy | open | hands | unspeeded |
| Heed et al. (2012) [23] - exp. 2 | 14 | $t$ | Accuracy | open | hands | unspeeded |
| Heed et al. (2016) [4] | 24 | $\beta$ | Accuracy | open | hands | NA |
| Kóbor et al. (2006) [17] | 18 | $F$ | JND 75% | closed | feet anatomical | unspeeded |
| Landry & Champoux (2018) [60] | 20 | $MD, SD_{within}$ | Accuracy | NA | feet external | unspeeded |
| Moseley et al. (2012) [11] | 10 | $t$ | JND 75% | open | verbal | NA |
| Nishikawa et al. (2015) [61] | 20 | $MD, SD_{within}$ | A | closed | hands | time restriction |
| Roberts & Humphreys (2008) [62] – exp. 1 | 9 | $t$ | slope | closed | feet external | unspeeded |
| Röder et al. (2004) [20] - eyes closed | 13 | $F$ | JND 75% | closed | hands | unspeeded |
| Röder et al. (2004) [20] - eyes open | 12 | $F$ | JND 75% | open | hands | unspeeded |
| Sambo et al. (2013) [63] | 12 | $t$ | JND 75% | closed | hands | time restriction |
| Schicke & Röder (2006) [27] | 10 | $F$ | JND 75% | closed | hands | unspeeded |
| Sharp et al. (2018) [64] | 13 | $MD, SD_{within}$ | Accuracy | NA | feet external | unspeeded |
| Shore et al. (2002) [2] | 9 | $F$ | slope | open | hands | unspeeded |
| Soto-Faraco & Azañón (2013) [10] | 32 | $F$ | JND 84% | open | hands | NA |
| Unwalla et al. (2020) [65] | 20 | $t$ | Accuracy | NA | hands | time restriction |
| Unwalla, Goldreich, et al. (2021) [15] – exp. 1 | 20 | $t$ | Accuracy | NA | feet anatomical | time restriction |
| Unwalla, Goldreich, et al. (2021) [15] - exp. 2 | 43 | $t$ | Accuracy | NA | feet anatomical | time restriction |
| Unwalla, Cadieux, et al. (2021) [18] - eyes closed | 20 | $t$ | Accuracy | closed | hands | time restriction |
| Unwalla, Cadieux, et al. (2021) [18] - eyes open | 20 | $t$ | Accuracy | open | hands | time restriction |
| Wada et al. (2004) [25] | 32 | $MD, SD_{within}$ | A | closed | hands | time restriction |

Note. Exp. = experiment, $n$ = simple size, $M$ = mean, $SE$ = standard error, $MD$ = mean difference, $SD$ = standard deviation, NA = not available

Seven studies comprised 2-4 experiments each (see Supplementary Table B2). Because these experiments involved different participants, we treat them as independent. For further information and limitations regarding the coding of subgroup analysis variables outcome measure, response modality, and task instructions about speed see Supplementary Tables B3, B4, and B5.

Our main analysis involved the data from all identified 29 studies with their 39 experiments. The estimated crossing effect, quantified as Cohen’s *d*_*z*_, was 1.45 (*t*(38) = 20.26, *p* < .001, 95% CI [1.3, 1.59]). By Cohen’s convention, this is a very large effect size (large when *d*_*z*_ >= 0.8) [66]. Fig. 2 presents experiment-level estimates. The estimated between-experiment heterogeneity variance, *τ*^2^, was 0.07 (95%CI [0.00, 0.15]), with Q (38) = 52.38, *p* = .06, and an *I*^2^ value of 28% (95%CI [0%, 51%]). The prediction interval ranged from Cohen’s *d*_*z*_ = 0.88 to 2.01, indicating that the TOJ crossing effect is large and should be reliably observable in future studies.

**Figure 2.**
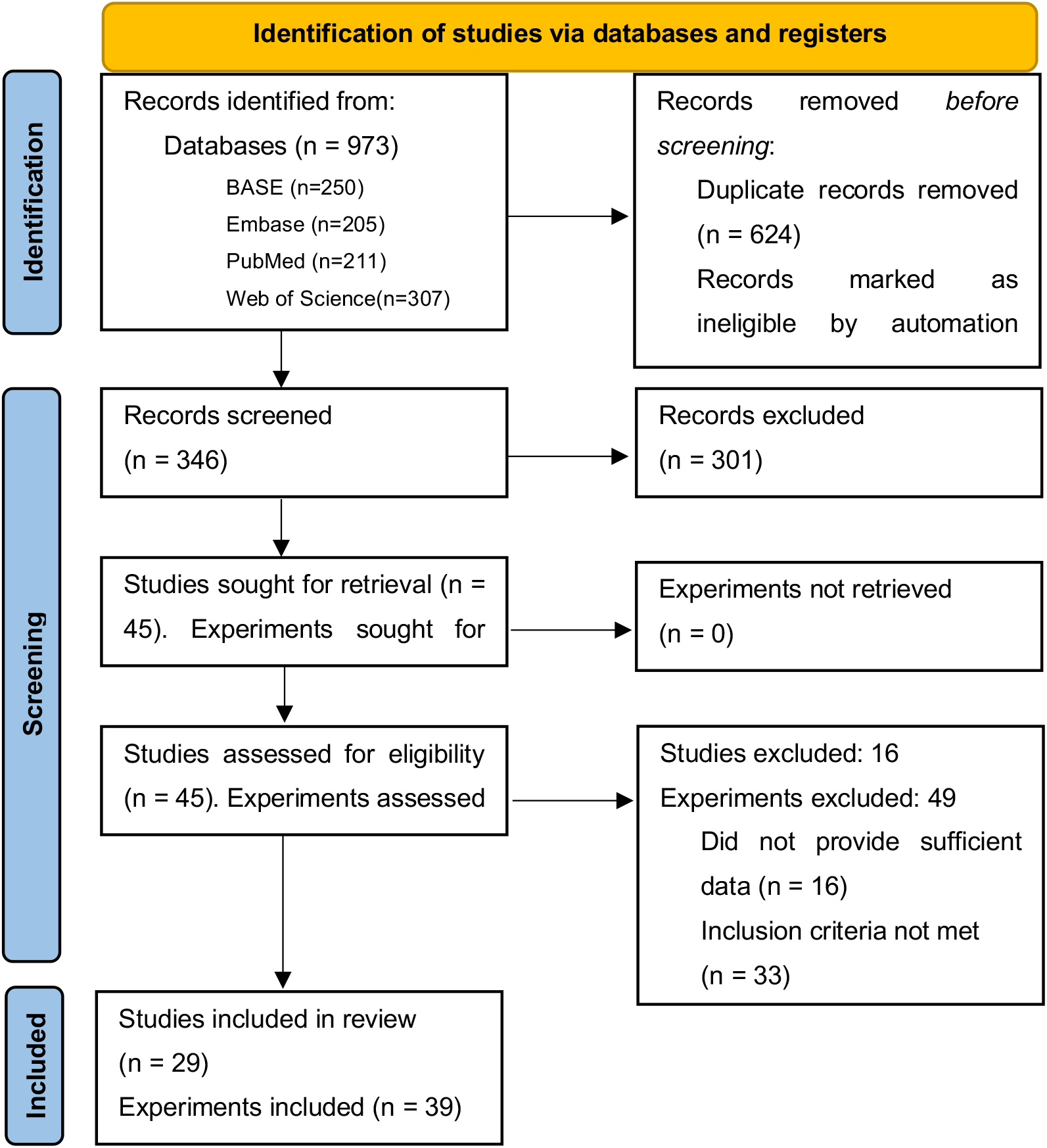
Flowchart of study selection. Design adapted from the PRISMA 2020 statement [51].

The outlier analysis identified two experiments: Cadieux et al. (2010; experiment 15 in Fig. 3) [21] and Soto-Faraco and Azanon (2013; experiment 32 in Fig. 3) [10]. The same two studies also contributed strongly to overall heterogeneity measured by Cochran’s Q and showed high influence in a leave-one-out analysis (see Fig. 3). The effect size in Cadieux et al. (2010) exceeded the pooled estimate, whereas the effect in Soto-Faraco & Azañón (2013) was smaller. Excluding these two experiments affected the numerical results only slightly (Cohen’s *d*_*z*_, = 1.43, *p* < .001, 95% CI [1.29, 1.57], 95% PI [1.1, 1.76; *I*^2^ value of 2%, 95%CI [0%, 39%]).

**Figure 3.**
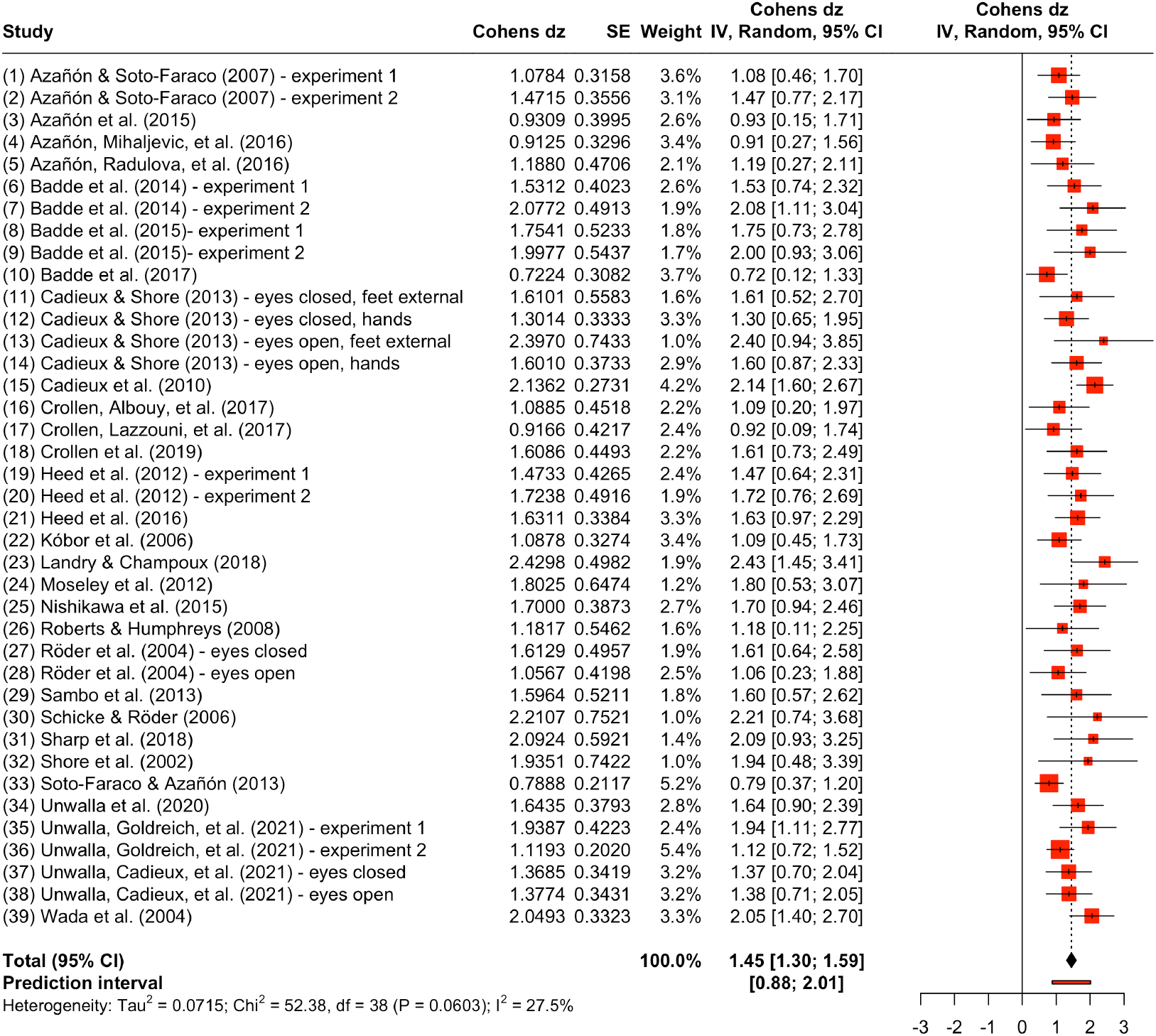
Forest plot of the effect sizes of included experiments. *SE* = standard error, CI = confidence interval, *df* = degrees of freedom. Red squares represent the point estimates for each experiment and black lines are 95% confidence intervals. The size of the squares reflects the study weight with which a study contributes to the pooled effect size. The black diamond symbolizes the pooled effect estimate, while the red line below denotes the prediction interval.

#### Influence of Moderators on Effect Size Estimate

Table 2 summarizes influence of the four potential moderators – analysis method, visual input, response mode, and speed and accuracy instructions – in the same order as above.

**Table 2.**
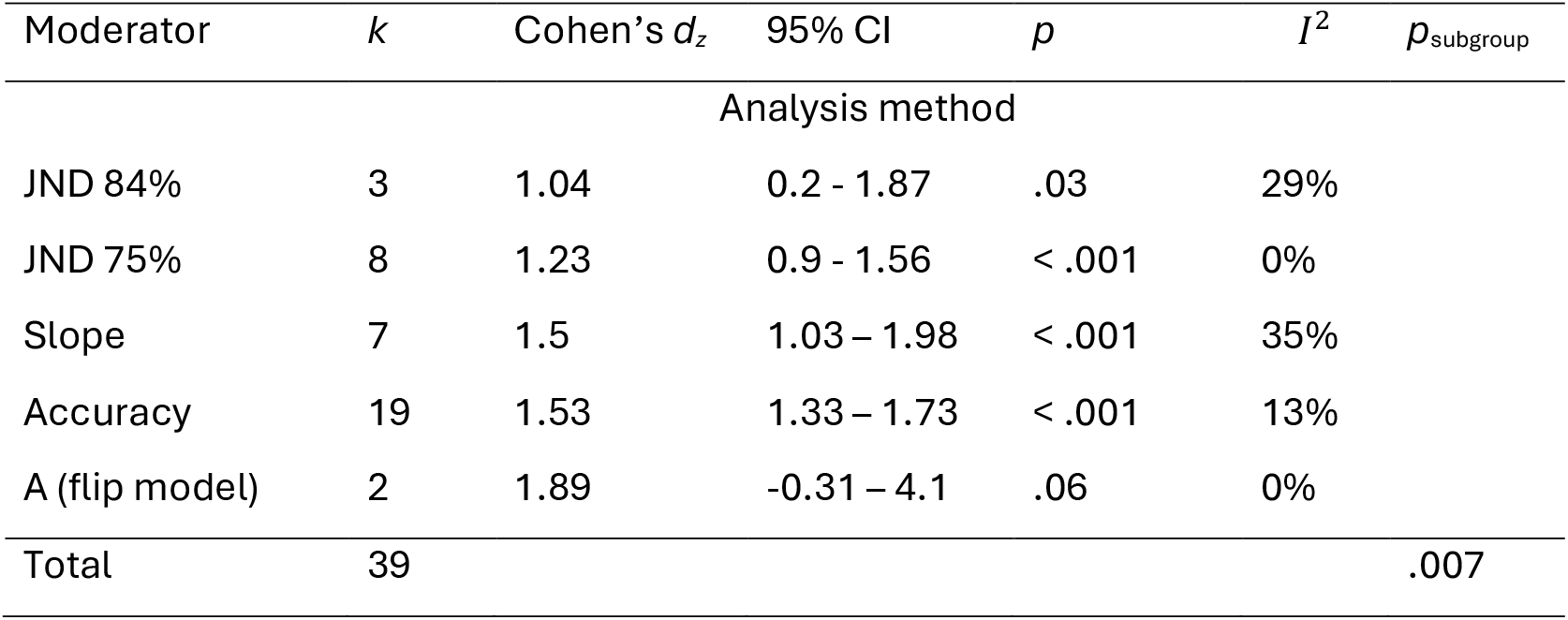
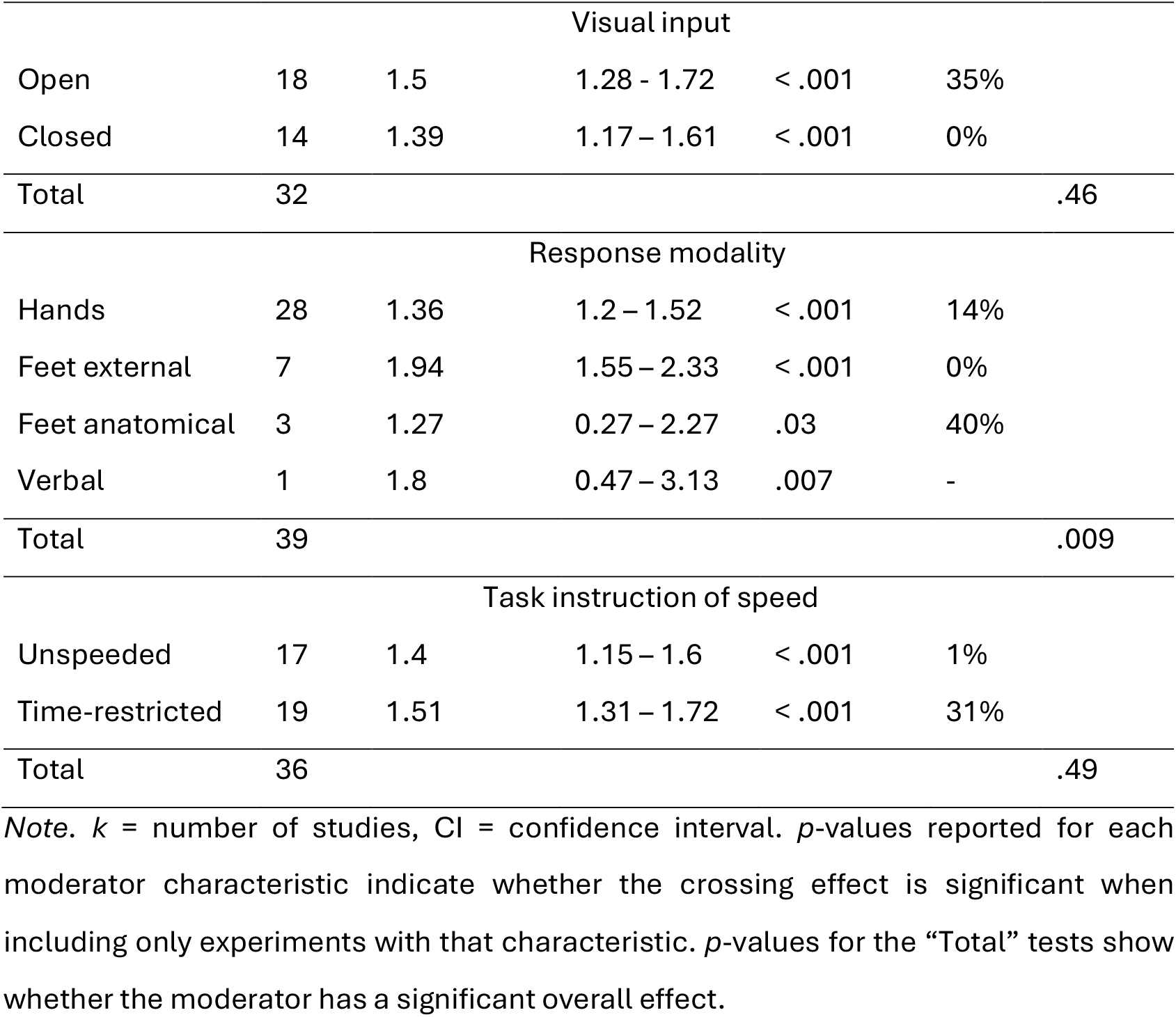
Effect size estimates when different potential moderators were considered.

For the analysis method, the largest descriptive crossing effects was present for slope, accuracy, and the flip model’s A parameter, a slightly smaller crossing effect for JND 75%, and the smallest for JND 84%. Nevertheless, all effect size estimates were large, ranging from 1.04-1.89. Differences between methods were significant overall (p = 0.007) e. Pairwise post-hoc comparisons are listed in Table 3.

**Table 3.** Post-hoc pairwise comparisons of the moderator analysis method using the Tukey test.

| Analysis Method | *estimate* | *se* | *t* | *df* | *p* |
| --- | --- | --- | --- | --- | --- |
| A – Accuracy | -0.37 | 0.28 | -1.28 | 34 | .208 |
| A – JND 75% | -0.66 | 0.31 | -2.13 | 34 | .041 |
| A – JND 84% | -0.86 | 0.33 | -2.62 | 34 | .013 |
| A – slope | -0.39 | 0.32 | -1.23 | 34 | .228 |
| Accuracy – JND 75% | -0.3 | 0.19 | -1.61 | 34 | .117 |
| Accuracy – JND 84% | -0.49 | 0.21 | -2.35 | 34 | .025 |
| Accuracy – slope | -0.02 | 0.19 | -0.12 | 34 | .904 |
| JND 75% – JND 84% | -0.19 | 0.25 | -0.79 | 34 | .437 |
| JND 75% – slope | 0.27 | 0.23 | 1.18 | 34 | .245 |
| JND 84% – slope | 0.47 | 0.25 | 1.86 | 34 | .072 |

Note. se = standard error, *df* = degrees of freedom.

For visual input, the crossing effect was descriptively larger with eyes open than with eyes closed. However, this difference was not statistically significant, and the effect size was very large in both cases.

For response modality, the crossing effect was smaller for foot responses under anatomical instructions and for hand responses, and larger for verbal responses and for foot responses under external-spatial instructions. Note, however, that only a single experiment employed verbal responses, and three experiments employed anatomical foot responses. Response methods differed significantly overall (p = 0.009). This overall effect was driven by significant differences between hand responses vs. external foot responses and anatomical vs. external foot responses (see Table 4).

**Table 4.** Post-hoc pairwise comparisons of the moderator response modality using the Tukey test.

| Response Modality | *estimate* | *se* | *t* | *df* | *p* |
| --- | --- | --- | --- | --- | --- |
| feet anatomical – feet external | 0.67 | 0.26 | 2.59 | 35 | .014 |
| feet anatomical – hands | 0.09 | 0.2 | 0.45 | 35 | .655 |
| feet anatomical – verbal | 0.53 | 0.65 | 0.82 | 35 | .417 |
| feet external – hands | -0.58 | 0.19 | -2.99 | 35 | .005 |
| feet external – verbal | -0.14 | 0.65 | -0.21 | 35 | .832 |
| hands – verbal | 0.44 | 0.63 | 0.71 | 35 | .485 |

Note. se = standard error, *df* = degrees of freedom.

The crossing effect was smaller for unspeeded than speeded responses, but this descriptive difference was not statistically significant.

#### Assessment of potential publication bias

Meta-analyses can suffer from publication bias: when studies with non-significant results are less likely to be published, effect sizes may be systematically overestimated. We assessed potential publication bias in three ways. First, a contour-enhanced funnel plot (Fig. 4A) showed a paucity of studies with small effects (low Cohen’s d_z_) at high variability (large standard errors). If publication were independent of statistical significance, the region within the dotted significance contours would be expected to be symmetrically populated. The relative absence of non-significant results in these regions is commonly interpreted as underrepresentation due to selective publication of significant findings.

**Figure 4.**
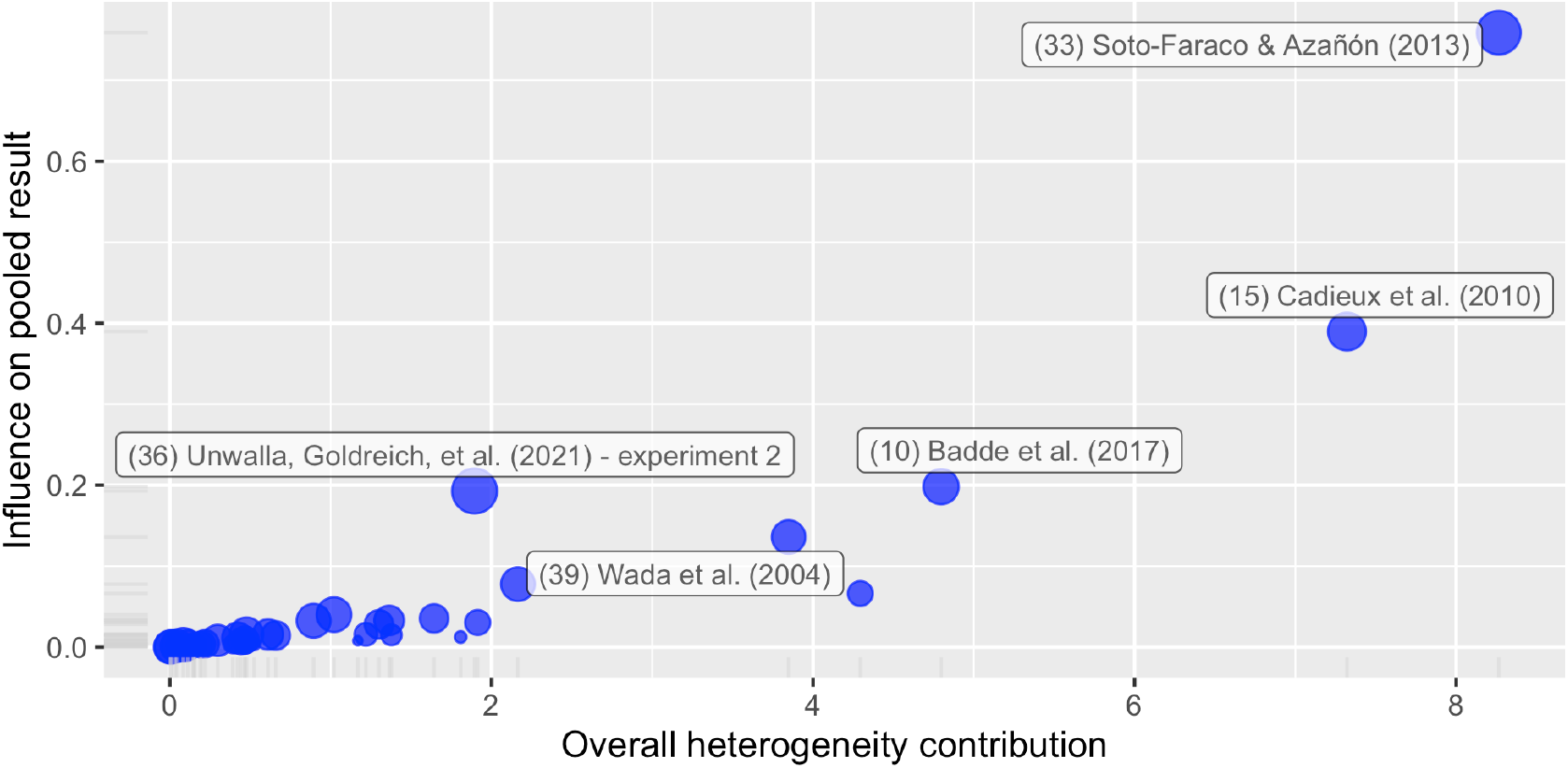
Baujat Plot. Each point represents an experiment included in the main meta-analysis, illustrating its contribution to heterogeneity and influence on the pooled effect; x-axis: contribution to heterogeneity (Cochran’s *Q*component); y-axis: influence on the pooled effect, quantified as the absolute change in the overall estimate from leave-one-out analysis (in units of Cohen’s *d*_*z*_). Only positive values are shown on the y-axis because both positive and negative deviations are plotted as absolute values. Circle size indicates the experiment’s weight in the meta-analysis. See main text for details.

**Figure 5.**
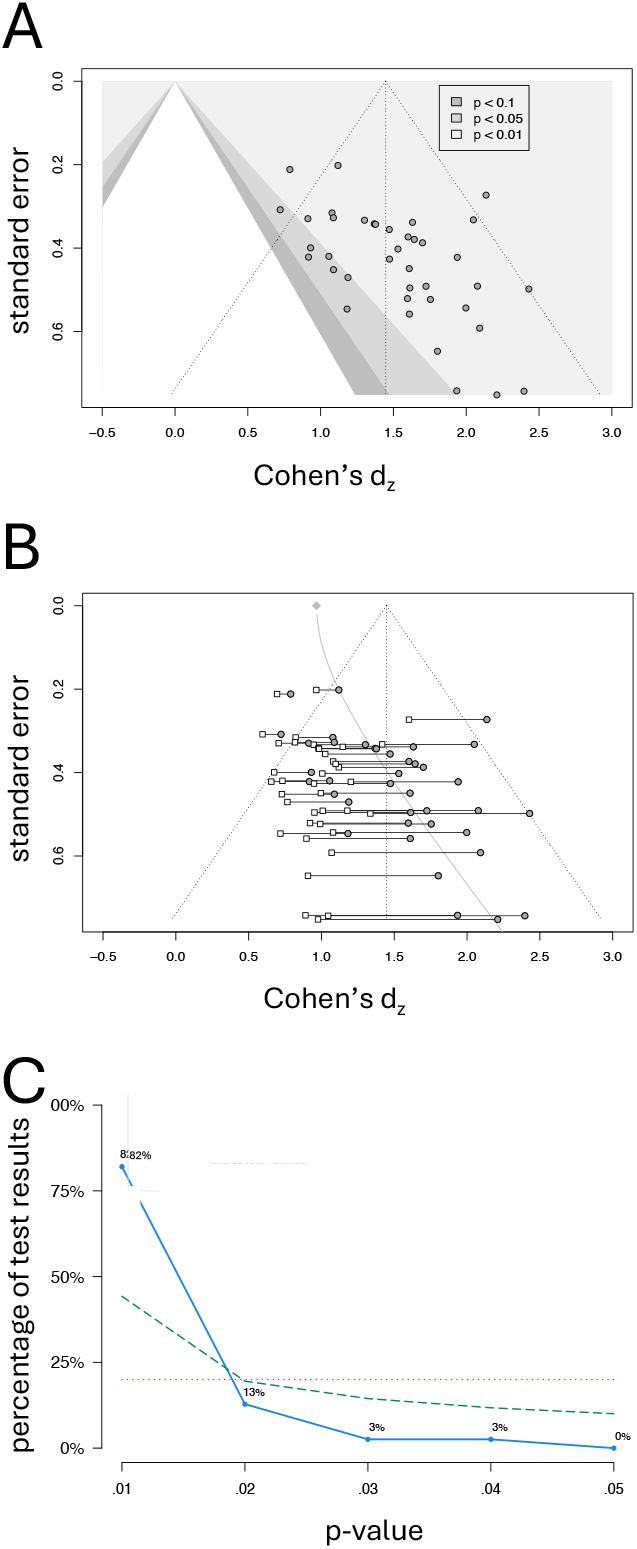
Assessment of potential publication bias. (A) Contour-enhanced funnel plot. Each point is an experiment’s effect size (Cohen’s d_z_) plotted against its standard error. The dashed vertical line marks the meta-analytic mean; the dashed triangle shows the ideal funnel shape expected if results are independent of statistical significance. The white funnel indicates the expected spread if no true effect existed; shaded contours denote p-value significance regions. (B) Funnel Plot with bias adjustment according to Rücker [50]. Original estimates (grey circles) and shrunken estimates (white squares) are linked for each experiment. The grey curve depicts the limiting adjusted mean effect (precision → ∞; standard error on the y-axis → 0), illustrating increasing small-study bias as standard error grows. (C) Observed p-curve (blue line): distribution of reported p-values. The red dotted line shows the null hypothesis (no true effect; uniform p-values), evaluated with the right-skewness test. The green dashed line shows a low-power scenario (small true effect; β = 0.33), evaluated with the flatness test.

Second, a funnel plot with Rücker-type bias adjustment, which models the relationship between effect size and precision to correct for small study effects, showed shrinkage towards an effect size of 0.97, compared to the original estimate of 1.45 (Fig. 4B). The larger the effect, the stronger was the correction, yielding a clustering of adjusted estimates around the corrected mean (see white dots in Fig. 4B). Taken together, the pattern observed in the two funnel-plot analyses is typically interpreted as evidence for small-study effects, potentially reflecting publication bias.

Therefore, we conducted a third test by examining the distribution of reported p-values (Fig. 4C). Specifically, we compared the observed distribution to that expected under the null hypothesis of no true effect and under a low-power scenario (β = 0.33). The observed distribution did not show an excess of p-values just below 0.05, as would be expected under p-hacking, and the distribution was also inconsistent with the absence of a true effect (right-skewness test: p < .001) and uniformly low power (flatness test: p > .999). The effect size estimate based on the p-curve was 1.4, a value almost identical to the original estimate of our meta-analysis, and power of the published experiments was estimated at β = 0.94.

In summary, these analyses indicate that the TOJ crossing effect is robust and large, although its magnitude in the published literature is likely somewhat overestimated. Bias assessment was less optimistic with funnel plots than with p-curve analysis, though the respective power estimates were always high (>0.8) in Cohen’s effect size classification.

## Discussion

Our meta-analysis over 29 studies comprising 39 experiments provides robust evidence for a significant crossing effect in tactile TOJ tasks–that is, a pronounced performance decline when the arms are crossed. The pooled effect size was Cohen’s *d*_*z*_ = 1.45, with a prediction interval from 0.88 to 2.01. Funnel-based correction lowered the mean effect to 0.97, and p-curve-based correction to 1.4. Thus, even the most conservative estimate classifies the effect as large. Between-study heterogeneity was low, further supporting the reliability of our assessment that the TOJ crossing effect is large. These formal results align with the field’s view of the TOJ crossing as a strong phenomenon that emerges even in small samples. The p-curve analysis supports this view, showing only a minimal reduction in the effect size estimate and indicating high power for published TOJ experiments. Nevertheless, the funnel analyses suggest that non-significant results may be underrepresented in the literature. It is important to note that small-study effect methods can be biased when the standard error depends on the effect size [67], as is the case here, and that p-curve analyses are accurate only when the analyzed p-value directly tests the target hypothesis [68], which is rarely the case in the TOJ literature. Overall, our bias assessment did not raise red flags that would cast doubt on the existence of a strong tactile TOJ crossing effect.

These claims pertain to the crossing effect per se, not to the moderator conditions we examined or to additional manipulations present in many source studies that we did not analyze. The crossing effect remained large across all moderator levels, indicating a robust phenomenon largely independent of assessment specifics. By contrast, moderator-related modulations were smaller. Reliable estimation of these smaller effects will require dedicated studies and substantially more replications. At present, the number of experiments is too small for precise moderator estimates; accordingly, our moderator analyses should be regarded as exploratory. Moreover, if publication bias exists in this literature, it is likely to have a greater impact on moderator effects (given their smaller magnitudes) than on the primary TOJ crossing effect.

Nevertheless, the effect size was affected by the analysis method. A previous report that examined three experiments with multiple analysis approaches [23], reported qualitatively comparable results across methods, and none showed systematic biases, such as being always more sensitive or always yielding larger effects than others. Here, the A parameter of the flip model exhibited the largest effect size. However, this estimate is based on only two studies, and exploratory data analysis of two of our own experiments suggested that the flip model’s A parameter produces effect sizes well in the range of the other measures (i.e., nearer to 1). Therefore, we do not interpret the present results as indicating superiority of the A parameter over other experimental measures.

Numerically, slope, accuracy, and flip measures produced larger crossing effects than JND measures. Incidentally, the JND has been criticized because it is computed as a ratio of the SOA required for stimulus identification and the estimated slope. This division introduces a non-linearity that can overemphasize poor performance (i.e., small slopes inflate the JND). Moreover, the characteristic N-shaped psychometric pattern with crossed hands can yield negative slope estimates for some participants, which is conceptually problematic and misrepresents the underlying response distribution. Consistent with these concerns, recent studies have tended to favor the alternative measures. In sum, our results, together with previous work suggest that analysis choices are probably not a major concern when comparing studies in this field.

Our moderator analysis of response mode suggests that the spatial content of response instructions affects the TOJ crossing effect, in line with reports from studies using both TOJ [69] and other paradigms [70,71]. The crossing effect was larger when foot responses were given under external instructions (i.e., “on which side of space did the first stimulus occur”), both against foot responses under anatomical instructions (i.e., “respond with the foot of the same body side as the hand stimulated first”) and against hand responses, for which instructions are typically anatomically formulated (“respond with the hand that received the first stimulus”). In sum, these findings suggest that TOJ performance is impaired by external-spatial response instructions, regardless of the responding effector.

Descriptively, the crossing effect was smaller when participants were blindfolded than when they could see. Although this difference did not reach statistical significance, its direction aligns with hypotheses about the role of visual information in tactile processing [20]. Notably, some prior studies have reported a strong visual influences [14]. The relatively small difference observed here may reflect our coarse grouping: we coded experiments only by whether the eyes were open or closed, without considering other factors, such as room illumination and fixation control. Because some tactile brain responses are modulated by gaze direction [72,73], there is a theoretical rationale to examine fixation. However, we are aware of only a single study that directly tested gaze effects on TOJ [4], and it reported a small effect. When gaze is not controlled, random variation of gaze behavior across trials and participants may limit systematic differences between experiments that do vs. do not endorse fixation. Given the limited power of our moderator analyses, we did not further subdivide experiments by fixation. Similarly, we know of only one study that systematically manipulated lighting [14], and it did not find effects of the light on vs. off. As most studies provide little detail on lighting conditions, this factor was not amenable to analysis here.

We observed a slight tendency toward a more pronounced crossing effect in experiments that imposed time constraints compared with those that did not. However, this difference was small and did not reach statistical significance. Extracting this information from the original reports proved challenging, and we could only apply a coarse “speeded vs. unspeeded” categorization. More detailed reporting in future studies would enable finer-grained analysis and may clarify whether time pressure amplifies the crossing effect.

In sum, even though some moderators of the TOJ crossing effect have received attention in research and show numerically suggestive patterns, the statistical picture remains inconclusive. Firmly establishing the role of these moderators will require additional experiments and diligent reporting, including non-significant findings. Moreover, these moderators are not merely task-level nuisance factors but are theoretically interesting in their own right. Accordingly, they could be fruitfully investigated across multiple paradigms, not only in TOJ.

Where can the field go from here? The overall picture is encouraging for using the TOJ crossing effect in research on tactile spatial processing. Indeed, several studies have leveraged this task to address questions beyond basic tactile research, including clinical and developmental contexts [8–11]. When an experimental effect is used as a marker (e.g., for a clinical condition or a developmental process), it should meet four criteria:

1. Magnitude. The effect should be large enough to detect with reasonable sample sizes. This criterion is clearly met, as shown here.
2. Reliability. The effect should be stable over time. Indeed, the TOJ task has been attested high reliability [65].
3. Between-participant variance. There should be sufficient variability to allow correlations with constructs relevant to the research question. For instance, one might test whether TOJ performance is impaired in individuals on the autism spectrum, based on the hypothesis that TOJ performance correlates with other aspects of sensorimotor processing [8] and autism may involve sensorimotor deficits. If between-participant variance were minimal, such correlations would be unlikely [74,75]. Fortunately, the TOJ task shows substantial interindividual variability (see [21] for single-participant illustrations).
4. Validity. The task should map onto the intended underlying construct. At present, adequate studies directly addressing the validity of TOJ are lacking. Tactile crossing effects occur not only in TOJ but also in other paradigms [6,76], and it is commonly assumed that these effects share a common tactile–spatial mechanism. However, dedicated evidence is scarce, and preliminary data from our laboratory challenge this assumption.

In our view, a priority in TOJ research is to establish the paradigm’s theoretical meaning by relating it to other variables and tasks. It is also noteworthy that both fine-grained investigations of (presumably smaller) moderator effects and correlational studies will require substantially larger samples than are typical in the TOJ—and more broadly in the sensorimotor and multisensory—literatures. Thus, although the TOJ crossing effect is large and detectable in small samples, answering broader mechanistic and translational questions will require larger, well-powered studies.

Beyond situating the TOJ crossing effect within a broader research scope, it will be important to further scrutinize its neural implementation. Prior studies using functional, resting-state, and structural MRI have implicated frontal and parietal cortices in the TOJ crossing effect [77–81], consistent with broader evidence for a central role of these regions in tactile–spatial processing [72,73,82,83]. Brain imaging research has also addressed theoretical questions, such as the role of vision in the development of tactile–spatial processing [77]. In this light, brain imaging—including EEG and MEG [10,73]–may be leveraged to probe the moderators discussed here, such as spatial instructions, response modality, and the availability of visual information. As just one example, imaging could help dissociate brain regions that contribute to TOJ generally from those recruited specifically under explicitly spatial instructions. It could also address underexplored questions about functional implementation, such as potential hemispheric lateralization of tactile–spatial functions, as suggested by differences in TOJ performance between left- and right-handers [25].

In sum, both behavioral and neural approaches have strong potential to (i) elucidate the mechanisms underlying the tactile TOJ crossing effect and (ii) harness the paradigm to address a broader range of theoretical and translational questions.

### Limitations

Our screening process was performed primarily by one author, so it is possible that we overlooked eligible studies. To mitigate this risk, we discussed inclusion whenever uncertainty arose, and we judge the likelihood of misclassification to be low. Nonetheless, some inclusion/exclusion decisions are inevitably subjective, and of the 85 experiments identified as principally eligible, we excluded 33 based on our selection criteria. We were also struck that about 20% of eligible studies (16 of 85) lacked sufficient statistics for meta-analysis (including, embarrassingly, some of our own). This attrition may bias our findings and may not fully reflect all published TOJ research. That said, we did not notice any obvious discrepancies between our meta-analytic results and the reported statistics of excluded studies.

We also found it unexpectedly difficult to extract several methodological details. Future work can facilitate meta-analytic efforts by thoroughly reporting experimental parameters—even those that may seem peripheral at the time of publication. This is especially pertinent because the TOJ paradigm is often used to test additional experimental manipulations rather than the crossing effect per se.

## Conclusion

In summary, we confirm that the tactile TOJ crossing effect is large, regardless of specific design aspects and analysis choices. The effect size estimate of approximately 1.4 may aid planning future studies, especially if they target the crossing effect per se, for instance, in clinical and developmental studies. By contrast, the evidence for moderators commonly discussed in the TOJ literature is far less conclusive; clarifying their roles will require additional research and thorough reporting. The same need applies to embedding the TOJ paradigm into a broader theoretical framework to establish its construct validity, an essential step if the task is to serve as a marker for clinically and developmentally relevant processes.

## Acknowledgments

We thank Tine-Marie Wujciak for helpful discussion during conceptualization, data curation, and formal analysis.

## Appendix A

**Table A1.** Search String for Each Database.

| Database | Search string |
| --- | --- |
| BASE | (tactile OR hand OR arm) temporal order judg* doctype:121;<br>english manually selected |
| EMBASE | ('tactile temporal order' OR ((hand OR tactile OR arm) AND<br>temporal AND order AND judg*)) AND ([article]/lim OR [article<br>in press]/lim) AND [english]/lim |
| PUBMED | (tactile OR hand OR arm) AND temporal order judg* AND<br>(english[Filter]) NOT(review pt]) |
| Web of Science | ALL=((hand OR tactile OR arm) temporal order judg*); articles<br>manually selected |

## Appendix B

Transformation of noncentral t variance to variance of Cohen’s d_z_

To calculate the standard error of Cohen’s *d*_*z*_, typically either the standard deviation of the mean difference or the correlation between the two measurements (crossed and uncrossed conditions) is needed [29,30]. However, these data were not available in all studies. Therefore, to avoid using an estimated correlation for the standard error calculation, the relationship between *t* and Cohen’s *d*_*z*_ was utilized to calculate the standard errors of the effect sizes:

1. Variance of the random variable t which follows a non-central t-distribution [84] (using an approximation [85,86]):

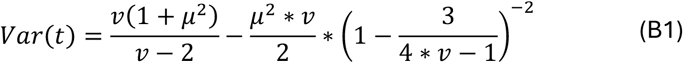

where *v* is the degrees of freedom and *μ* is the noncentrality parameter.
2. Given the relation between t and Cohen’s d_z_ [30], *t* can be calculated from *d*_*z*_, and the sample size:

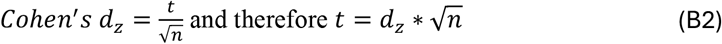
3. Substitution of *v* and *μ* for within-subject designs: where *v* = n-1 and 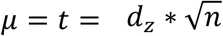

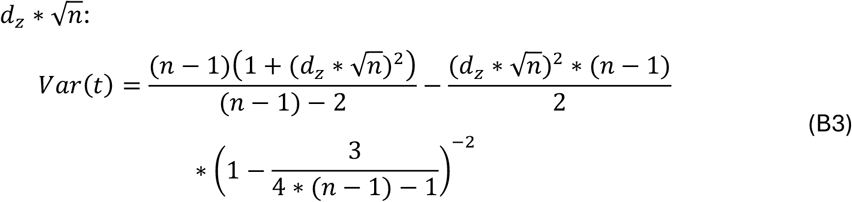
4. Given equation (B2) the relationship between *Var*(*t*) and *Var*(*d*_*z*_) can be expressed as:

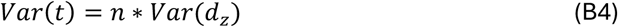

therefore:

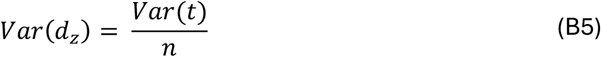
5. Substitution of *Var*(*t*) into the equation for *Var*(*d*_*z*_):

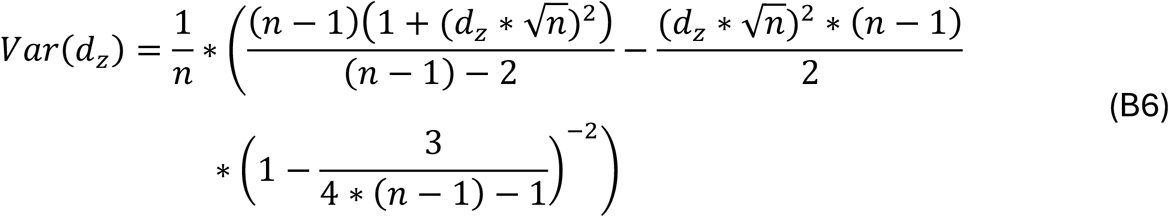
6. Simplified to:

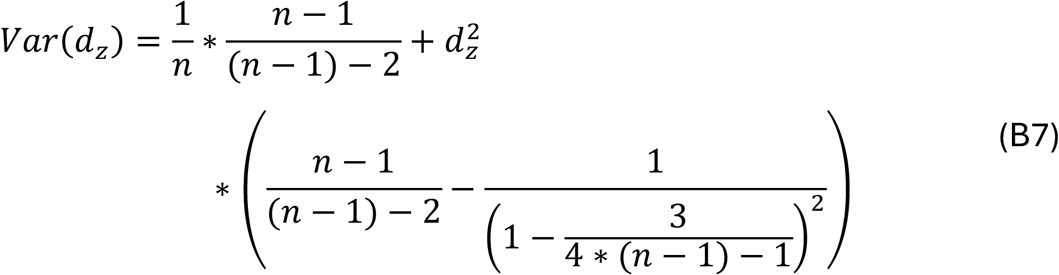
7. Derive the standard error SE(*d*_*z*_), which in this case equals to the standard deviation and is therefore defined as the square root of the variance:

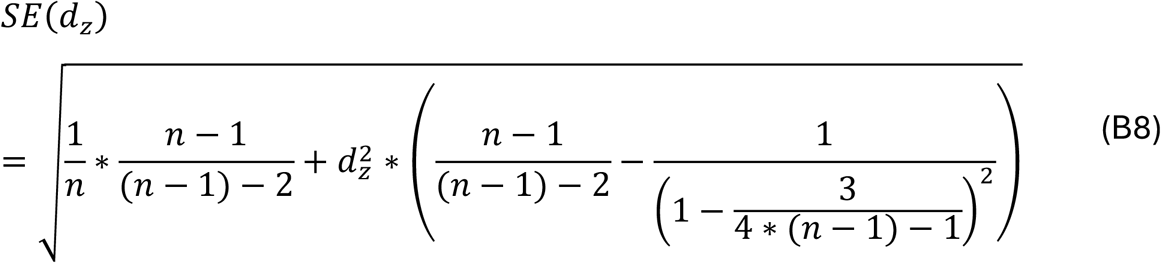

