## supplement 1 for "Crossing effects in the tactile temporal order judgment task: A meta-analysis"

### Supplementary Information for “Crossing effects in the tactile temporal order judgment task: A meta-analysis”

#### Supplement A

Correlation of uncrossed and crossed conditions

The correlation of uncrossed and crossed conditions was required for our analysis but was not reported in any study. Using a default correlation of .5 may lead to inaccuracies by overestimating the effect size if the true correlation exceeds .5 and by underestimating it if the true correlation is less than .5. We computed the correlation by utilizing openly accessible raw data obtained from studies that were part of the present meta-analysis. We accessed data from the following studies: Azañón, Mihaljevic, et al. (2016)[1], Heed et al. (2016)[2], Sharp et al. (2018)[3]. We selected subsets of the available data that closely aligned with a typical TOJ task to estimate the crossing effect (Table A1). Afterwards, we calculated a mean for the uncrossed and crossed conditions per participant and merged all data into one dataset. From this dataset the correlation between both conditions was calculated, $r(51) = .52$, $p < .001$, 95% CI [.29, .69].

**Table A1. Data subset used to calculate the correlation between uncrossed and crossed conditions**

| Study | Data subset |
|---|---|
| Azañón, Mihaljevic, et al. (2016)[1] | Condition “Aligned” |
| Heed et al. (2016)[2] | Condition “All-straight” |
| Sharp et al. (2018)[3] | Control group |

However, it is important to highlight that the correlation between both conditions varied strongly between studies, as shown in Table A2.

**Table A2. Correlation coefficients for each study**

| Study | *r* | df | 95% CI | *p* |
|---|---|---|---|---|
| Azañón, Mihaljevic, et al. (2016)[1] | .69 | 14 | .29 - .88 | .003 |
| Heed et al. (2016)[2] | .09 | 22 | -.32 - .48 | .675 |
| Sharp et al. (2018)[3] | .5 | 11 | -.07 - .82 | .083 |

Note. CI = confidence interval, *df* = degrees of freedom.

Furthermore, combining all the data into a single dataset assumes data independence and overlooks potential dependencies within each study. To address these limitations, we assessed the validity of the estimated correlation in two ways: First, replacing the estimate by extreme values (0.001; 0.999) in Equation 6 changed effect size estimates only minimally (Table A3). Second, computing Cohen's $d_z$ with Equation 6 including only those studies that had reported effect sizes, *t*-, or *F*-values, too, led to only a small modulation of effect size estimates (Table A4).

**Table A3. Effect sizes (Cohen's $d_z$) for different correlation estimates.**

| | Cohens $d_z$ calculated from | | | | |
|---|---|---|---|---|---|
| Study | *r* = .001 | *r* = .09 | *r* = .518 | *r* = .69 | *r* = .999 |
| Azañón & Soto-Faraco (2007)[4] – exp. 1 | 1.02 | 1.03 | 1.08 | 1.1 | 1.14 |
| Azañón & Soto-Faraco (2007)[4] – exp. 2 | 1.33 | 1.35 | 1.47 | 1.53 | 1.65 |

*Note.* Exp. = experiment

**Table A4. Effect size (Cohen's $d_z$) comparisons for studies where t-/F-value and mean and standard deviations of the same measurement were available.**

| | Cohens $d_z$ calculated from | | | | | |
|---|---|---|---|---|---|---|
| Study | *t*-/*F*-value | *r* = .001 | *r* = .09 | *r* = .518 | *r* = .69 | *r* = .999 |
| Cadieux et al. (2010)[5] | 2.14 | 1.88 | 1.95 | 2.5 | 2.89 | 4.69 |
| Crollen, Albouy, et al. (2017)[6] | 1.09 | 0.94 | 0.96 | 1.13 | 1.22 | 1.48 |

| | | | | | | |
|---|---|---|---|---|---|---|
| Crollen, Lazzouni, et al. (2017)[7] | 0.92 | 0.93 | 0.93 | 0.97 | 0.98 | 1.01 |
| Heed et al. (2012)[8] | 1.72 | 1.7 | 1.76 | 2.17 | 2.45 | 3.38 |
| Moseley et al. (2012)[9] | 1.8 | 1.39 | 1.46 | 2 | 2.49 | 20.16 |

### Supplement B

**Table B1. Experiments That Did Not Meet the Inclusion Criteria: References and Reasons for Exclusion.**

| Study | Reason for exclusion |
|---|---|
| Azañón et al. (2010)[10] - exp. 1 - 3 | Dependent Variable = Visual performance |
| Azañón et al. (2018)[11] | Mean age = 8.24 |
| Azañón et al. (2015)[12] - exp. 2 & 3 | Did not provide sufficient data (no main effect of crossing status reported) |
| Azañón et al (2015)[12] – exp. 4 | Visual-tactile task |
| Azañón, Mihaljevic, et al. (2016)[1]– exp. 1 & 2 | Hands were vertically misaligned |
| Azañón, Radulova, et al. (2016) [13] – exp. 1 & 2 | Stimulation on shoulders and elbows |
| Azañón, Radulova, et al. (2016) [13] – exp. 3 | Stimulation on a single skin location |
| Badde et al. (2017)[14] – exp. 2 | Stimulation with multiple stimuli |
| Cadieux & Shore (2013)[15] – Eyes open, lights off | Eyes open, but light off |
| Cadieux & Shore (2013)[15] – exp. 1b | Eyes open/closed as within-subject factor |
| Cadieux & Shore (2013)[15] – exp. 2 | Eyes open/closed as within-subject factor |
| Cadieux et al. (2010)[5] – exp. 2 | No TOJ-task |
| Craig & Belser (2006)[16] – exp. 1 & 2 | Did not provide sufficient data (only TOJ-data with training) |
| Craig & Belser (2006)[16] – exp. 3 | Did not provide sufficient data (only comparison between control group and musicians) |
| Crollen, Albouy, et al. (2017)[6] – exp. 2 | No TOJ-task |
| Di Pino et al (2020)[17] | No healthy control group |
| Ferri et al. (2016)[18] | Did not provide sufficient data (only time window of judgement reversal) |
| Heed et al. (2012)[8] – exp. 3 | Crossing of fingers within one hand |
| Heed et al. (2016)[2] – exp. 2 | Did not provide sufficient data (no main effect of crossing status reported) |
| Hense et al. (2019)[19] – exp. 1 | Did not provide sufficient data (no data from the control group) |
| Hense et al. (2019)[19] – exp. 2 | No TOJ-task |
| Manfron et al. (2021)[20] | No investigation of the crossing effect |
| Moharramipour & Kitazawa (2021)[21] | Did not provide sufficient data (no main effect of the crossing status) |
| Moharramipour et al. (2023)[22] – exp. 1 & 2 | Did not provide sufficient data (no behavioral data) |
| Moseley et al. (2012)[9] – exp. 2 - 5 | No healthy control group |
| Ora et al. (2016)[24] | Did not provide sufficient data (no behavioral data) |

| | |
|---|---|
| Ritterband-Rosenbaum et al. (2014)[25] – exp. 1 - 5 | Degrees of freedom do not fit number of participants |
| Roerts & Humphreys (2008)[26] – exp. 2 & 3 | No „typical“ TOJ-task |
| Shore et al. (2002)[27] – exp. 2 | Visual stimuli |
| Shore et al. (2002)[27] – exp. 3 | Same participants as in experiment 1 |
| Takahashi & Kitazawa (2007)[28] | Did not provide sufficient data (only graphical report of behavioral data) |
| Tamè et al. (2017)[29] | Localization task of fingers of one hand |
| Unwalla et al. (2020)[30] – exp. 2 | Did not provide sufficient data (only graphical report of crossing effect) |
| Wada et al. (2012)[32] | Did not provide sufficient data (only graphical report of behavioral data) |
| Wada et al. (2014)[33] | Mean age = 11.7 |
| Yamamoto & Kitazawa (2001)[34] | Did not provide sufficient data (no standard deviation of flip parameters) |

*Note*. Exp. = experiment

**Table B2. Limitations of included studies.**

| Author | Limitation | Dependency |
|---|---|---|
| Azañón & Soto-Faraco (2007)[4] - exp. 1 | Participants saw rubber hands (in the same position as their own hands) | Yes |
| Azañón & Soto-Faraco (2007)[4] - exp. 2 | Participants saw rubber hands (in the same position as their own hands) | Yes |
| Azañón et al (2015)[12] | Main effect (posture change block wise & after 1 – 3 trials) | No |
| Azañón, Mihaljevic, et al. (2016)[1] | - | No |
| Azañón, Radulova, et al. (2016)[13] | - | No |
| Badde et al. (2014)[35] - exp. 1 | Main effect (working memory load: no, low & high) | Yes |
| Badde et al. (2014)[35] - exp. 2 | Main effect (working memory load: no, low & high) | Yes |
| Badde et al. (2015)[36] - exp. 1 | Different stimulus frequency | Yes |
| Badde et al. (2015)[36] - exp. 2 | Different stimulus frequency | Yes |
| Badde et al. (2017)[14] | Main effect (normal & with sound in between the stimuli) | No |
| Cadieux & Shore (2013)[15] - eyes closed, feet external | - | Yes |
| Cadieux & Shore (2013)[15] - eyes closed, hands | - | Yes |
| Cadieux & Shore (2013)[15] - eyes open, feet external | - | Yes |
| Cadieux & Shore (2013)[15] - eyes open, hands | - | Yes |
| Cadieux et al. (2010)[5] | - | No |
| Crollen, Albouy, et al. (2017)[6] | - | No |
| Crollen, Lazzouni, et al. (2017)[7] | Participants completed the task while lying | No |
| Crollen et al. (2019)[37] | - | No |
| Heed et al. (2012)[8] - exp. 1 | Main effect (control, hands flat & turned) | Yes |
| Heed et al. (2012)[8] - exp. 2 | - | Yes |
| Heed et al. (2016)[2] | Main effect (all straight, eyes to side & eyes and arms to side) | No |
| Kóbor et al., 2006[38] | main effect (hands in front & behind the back) | No |
| Landry & Champoux (2018)[39] | - | No |
| Moseley et al. (2012)[9] | - | No |

| | | |
|---|---|---|
| Nishikawa et al. (2015)[23] | Values for the measure of variability were assumed to be standard deviations | No |
| Roberts & Humphreys (2008)[26] | Different stimulation frequency | No |
| Röder et al. (2004)[40] - eyes closed | - | Yes |
| Röder et al. (2004)[40] - eyes open | - | Yes |
| Sambo et al. (2013)[41] | - | No |
| Schicke & Röder (2006)[42] | Main effect (stimulation on hand & feet) | No |
| Sharp et al. (2018)[3] | - | No |
| Shore et al. (2002)[27] | - | No |
| Soto-Faraco & Azañón, 2013[43] | - | No |
| Unwalla et al. (2020)[30] | - | No |
| Unwalla, Goldreich, et al. (2021)[44] - exp. 1 | - | Yes |
| Unwalla, Goldreich, et al. (2021)[44] - exp. 2 | - | Yes |
| Unwalla, Cadieux, et al. (2021)[45] - eyes closed | - | Yes |
| Unwalla, Cadieux, et al. (2021)[45] - eyes open | - | Yes |
| Wada et al. (2004)[31] | Right- and left-handed participants | No |

*Note.* Exp. = experiment

**Table B3. Specifications for subgroup analysis outcome measure.**

| Study | Specification |
|---|---|
| Azañón et al. (2015)[12] | Logistic function to calculate JND 75% |
| Azañón, Mihaljevic, et al. (2016)[1] | Logit transformation |
| Azañón, Radulova, et al. (2016)[13] | Probit transformation |
| Badde et al. (2017)[14] | χ2-value from GLMM coded as accuracy |
| Crollen et al. (2019)[37] | Probit transformation |
| Crollen, Albouy, et al. (2017)[6] | Probit transformation |
| Crollen, Lazzouni, et al. (2017)[7] | Probit transformation |
| Kóbor et al. (2006)[38] | Weibull transformation |
| Moseley et al. (2012)[9] | Probit transformation |
| Roberts & Humphreys (2008)[26] | Logistic slope comparison without transformation |

| | |
|---|---|
| Röder et al. (2004)[40] - eyes closed | Probit transformation |
| Röder et al. (2004)[40] - eyes open | Probit transformation |
| Sambo et al. (2013)[41] | Not transformed Gaussian Cumulative Function |
| Schicke & Röder (2006)[42] | Probit transformation |

Table B4 Specifications for subgroup analysis response modality

| Study | Specification |
|---|---|
| Roberts & Humphreys (2008)[26] | Responses with a single foot (by lifting the toe or the heel) |
| Schicke & Röder (2006)[42] | Responses with hands and feet but coded as hands since participants were instructed to respond with the stimulated body part |
| Unwalla, Goldreich, et al. (2021)[44] - exp. 1 | Selection of Feet anatomical due to the limited availability of studies with that condition |
| Unwalla, Goldreich, et al. (2021)[44] - exp. 2 | Selection of Feet anatomical due to the limited availability of studies with that condition |

*Note.* Exp. = experiment

Table B5 Specifications for subgroup analysis task instructions about speed

| Study | Specification |
|---|---|
| Roberts & Humphreys (2008)[26] | Coded as unspeeded since accuracy was stressed rather than the speed of responding |
| Sambo et al. (2013)[41] | Coded as time restricted since participants were instructed to respond "as accurately and as rapidly as possible“ |

### Supplement C

#### Forest Plots grouped by moderator variable

| Study or Subgroup | Cohens dz | SE | Weight | Cohens dz IV, Random, 95% CI |
|---|---|---|---|---|
| **Analysis.method = JND 84%** | | | | |
| (1) Azañón & Soto-Faraco (2007) - experiment 1 | 1.0784 | 0.3158 | 3.6% | 1.08 [ 0.46; 1.70] |
| (2) Azañón & Soto-Faraco (2007) - experiment 2 | 1.4715 | 0.3556 | 3.1% | 1.47 [ 0.77; 2.17] |
| (33) Soto-Faraco & Azañón (2013) | 0.7888 | 0.2117 | 5.2% | 0.79 [ 0.37; 1.20] |
| **Total (95% CI)** | | | **11.9%** | **1.04 [ 0.20; 1.87]** |
| Heterogeneity: Tau$^2$ = 0.0430; Chi$^2$ = 2.81, df = 2 (P = 0.2448); I$^2$ = 28.9% | | | | |
| **Analysis.method = JND 75%** | | | | |
| (22) Kóbor et al. (2006) | 1.0878 | 0.3274 | 3.4% | 1.09 [ 0.45; 1.73] |
| (24) Moseley et al. (2012) | 1.8025 | 0.6474 | 1.2% | 1.80 [ 0.53; 3.07] |
| (27) Röder et al. (2004) - eyes closed | 1.6129 | 0.4957 | 1.9% | 1.61 [ 0.64; 2.58] |
| (28) Röder et al. (2004) - eyes open | 1.0567 | 0.4198 | 2.5% | 1.06 [ 0.23; 1.88] |
| (29) Sambo et al. (2013) | 1.5964 | 0.5211 | 1.8% | 1.60 [ 0.57; 2.62] |
| (3) Azañón et al. (2015) | 0.9309 | 0.3995 | 2.6% | 0.93 [ 0.15; 1.71] |
| (30) Schicke & Röder (2006) | 2.2107 | 0.7521 | 1.0% | 2.21 [ 0.74; 3.68] |
| (4) Azañón, Mihaljevic, et al. (2016) | 0.9125 | 0.3296 | 3.4% | 0.91 [ 0.27; 1.56] |
| **Total (95% CI)** | | | **17.8%** | **1.23 [ 0.90; 1.56]** |
| Heterogeneity: Tau$^2$ = 0.0430; Chi$^2$ = 5.39, df = 7 (P = 0.6122); I$^2$ = 0% | | | | |
| **Analysis.method = slope** | | | | |
| (15) Cadieux et al. (2010) | 2.1362 | 0.2731 | 4.2% | 2.14 [ 1.60; 2.67] |
| (16) Crollen, Albouy, et al. (2017) | 1.0885 | 0.4518 | 2.2% | 1.09 [ 0.20; 1.97] |
| (17) Crollen, Lazzouni, et al. (2017) | 0.9166 | 0.4217 | 2.4% | 0.92 [ 0.09; 1.74] |
| (18) Crollen et al. (2019) | 1.6086 | 0.4493 | 2.2% | 1.61 [ 0.73; 2.49] |
| (26) Roberts & Humphreys (2008) | 1.1817 | 0.5462 | 1.6% | 1.18 [ 0.11; 2.25] |
| (32) Shore et al. (2002) | 1.9351 | 0.7422 | 1.0% | 1.94 [ 0.48; 3.39] |
| (5) Azañón, Radulova, et al. (2016) | 1.1880 | 0.4706 | 2.1% | 1.19 [ 0.27; 2.11] |
| **Total (95% CI)** | | | **15.7%** | **1.51 [ 1.03; 1.98]** |
| Heterogeneity: Tau$^2$ = 0.0430; Chi$^2$ = 9.24, df = 6 (P = 0.1605); I$^2$ = 35.1% | | | | |
| **Analysis.method = Accuracy** | | | | |
| (10) Badde et al. (2017) | 0.7224 | 0.3082 | 3.7% | 0.72 [ 0.12; 1.33] |
| (11) Cadieux & Shore (2013) - eyes closed, feet external | 1.6101 | 0.5583 | 1.6% | 1.61 [ 0.52; 2.70] |
| (12) Cadieux & Shore (2013) - eyes closed, hands | 1.3014 | 0.3333 | 3.3% | 1.30 [ 0.65; 1.95] |
| (13) Cadieux & Shore (2013) - eyes open, feet external | 2.3970 | 0.7433 | 1.0% | 2.40 [ 0.94; 3.85] |
| (14) Cadieux & Shore (2013) - eyes open, hands | 1.6010 | 0.3733 | 2.9% | 1.60 [ 0.87; 2.33] |
| (19) Heed et al. (2012) - experiment 1 | 1.4733 | 0.4265 | 2.4% | 1.47 [ 0.64; 2.31] |
| (20) Heed et al. (2012) - experiment 2 | 1.7238 | 0.4916 | 1.9% | 1.72 [ 0.76; 2.69] |
| (21) Heed et al. (2016) | 1.6311 | 0.3384 | 3.3% | 1.63 [ 0.97; 2.29] |
| (23) Landry & Champoux (2018) | 2.4298 | 0.4982 | 1.9% | 2.43 [ 1.45; 3.41] |
| (31) Sharp et al. (2018) | 2.0924 | 0.5921 | 1.4% | 2.09 [ 0.93; 3.25] |
| (34) Unwalla et al. (2020) | 1.6435 | 0.3793 | 2.8% | 1.64 [ 0.90; 2.39] |
| (35) Unwalla, Goldreich, et al. (2021) - experiment 1 | 1.9387 | 0.4223 | 2.4% | 1.94 [ 1.11; 2.77] |
| (36) Unwalla, Goldreich, et al. (2021) - experiment 2 | 1.1193 | 0.2020 | 5.4% | 1.12 [ 0.72; 1.52] |
| (37) Unwalla, Cadieux, et al. (2021) - eyes closed | 1.3685 | 0.3419 | 3.2% | 1.37 [ 0.70; 2.04] |
| (38) Unwalla, Cadieux, et al. (2021) - eyes open | 1.3774 | 0.3431 | 3.2% | 1.38 [ 0.71; 2.05] |
| (6) Badde et al. (2014) - experiment 1 | 1.5312 | 0.4023 | 2.6% | 1.53 [ 0.74; 2.32] |
| (7) Badde et al. (2014) - experiment 2 | 2.0772 | 0.4913 | 1.9% | 2.08 [ 1.11; 3.04] |
| (8) Badde et al. (2015)- experiment 1 | 1.7541 | 0.5233 | 1.8% | 1.75 [ 0.73; 2.78] |
| (9) Badde et al. (2015)- experiment 2 | 1.9977 | 0.5437 | 1.7% | 2.00 [ 0.93; 3.06] |
| **Total (95% CI)** | | | **48.5%** | **1.53 [ 1.33; 1.73]** |
| Heterogeneity: Tau$^2$ = 0.0430; Chi$^2$ = 20.58, df = 18 (P = 0.3009); I$^2$ = 12.6% | | | | |
| **Analysis.method = A** | | | | |
| (25) Nishikawa et al. (2015) | 1.7000 | 0.3873 | 2.7% | 1.70 [ 0.94; 2.46] |
| (39) Wada et al. (2004) | 2.0493 | 0.3323 | 3.3% | 2.05 [ 1.40; 2.70] |
| **Total (95% CI)** | | | **6.1%** | **1.89 [-0.31; 4.10]** |
| Heterogeneity: Tau$^2$ = 0.0430; Chi$^2$ = 0.47, df = 1 (P = 0.4937); I$^2$ = 0% | | | | |
| **Total (95% CI)** | | | **100.0%** | **1.45 [ 1.30; 1.59]** |
| **Prediction interval** | | | | **[ 0.88; 2.01]** |

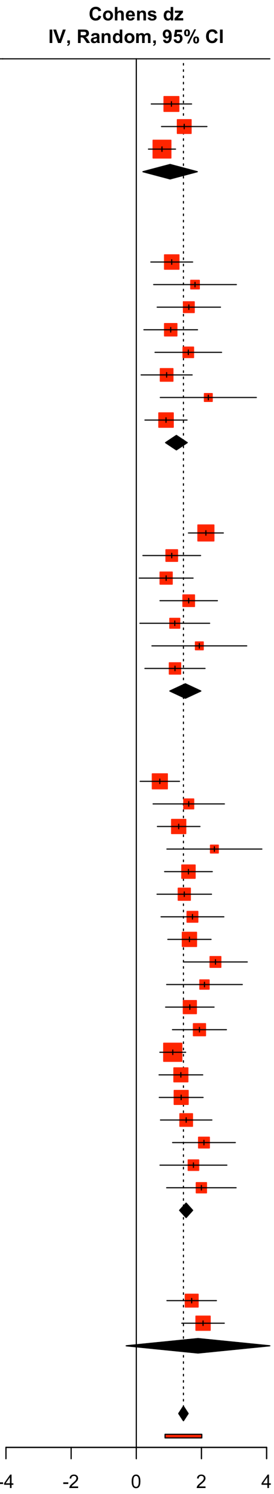


Heterogeneity: Tau$^2$ = 0.0715; Chi$^2$ = 52.38, df = 38 (P = 0.0603); I$^2$ = 27.5%
Residual heterogeneity: Tau$^2$ = 0.0430; Chi$^2$ = 38.50, df = 34 (P = 0.2731); I$^2$ = 11.7%
Test for subgroup differences: Chi$^2$ = 14.24, df = 4 (P = 0.0066)

**Figure C1. Forest plot of the effect sizes of included studies grouped by the moderator analysis method.** *SE* = standard error, CI = confidence interval, *df* = degrees of freedom. Red squares represent the point estimates for each study and black lines are 95% confidence intervals. The size of the squares reflects the study weight with which a study contributes to the pooled effect size. The black diamond symbolizes the pooled effect estimate, while the red line below denotes the prediction interval.

| Study or Subgroup | Cohens dz | SE | Weight | Cohens dz IV, Random, 95% CI |
|---|---|---|---|---|
| **Eyes = open** | | | | |
| (1) Azañón & Soto-Faraco (2007) - experiment 1 | 1.0784 | 0.3158 | 4.5% | 1.08 [0.46; 1.70] |
| (13) Cadieux & Shore (2013) - eyes open, feet external | 2.3970 | 0.7433 | 1.2% | 2.40 [0.94; 3.85] |
| (14) Cadieux & Shore (2013) - eyes open, hands | 1.6010 | 0.3733 | 3.6% | 1.60 [0.87; 2.33] |
| (15) Cadieux et al. (2010) | 2.1362 | 0.2731 | 5.3% | 2.14 [1.60; 2.67] |
| (19) Heed et al. (2012) - experiment 1 | 1.4733 | 0.4265 | 3.0% | 1.47 [0.64; 2.31] |
| (2) Azañón & Soto-Faraco (2007) - experiment 2 | 1.4715 | 0.3556 | 3.8% | 1.47 [0.77; 2.17] |
| (20) Heed et al. (2012) - experiment 2 | 1.7238 | 0.4916 | 2.4% | 1.72 [0.76; 2.69] |
| (21) Heed et al. (2016) | 1.6311 | 0.3384 | 4.1% | 1.63 [0.97; 2.29] |
| (24) Moseley et al. (2012) | 1.8025 | 0.6474 | 1.5% | 1.80 [0.53; 3.07] |
| (28) Röder et al. (2004) - eyes open | 1.0567 | 0.4198 | 3.0% | 1.06 [0.23; 1.88] |
| (3) Azañón et al. (2015) | 0.9309 | 0.3995 | 3.3% | 0.93 [0.15; 1.71] |
| (32) Shore et al. (2002) | 1.9351 | 0.7422 | 1.2% | 1.94 [0.48; 3.39] |
| (33) Soto-Faraco & Azañón (2013) | 0.7888 | 0.2117 | 6.9% | 0.79 [0.37; 1.20] |
| (38) Unwalla, Cadieux, et al. (2021) - eyes open | 1.3774 | 0.3431 | 4.0% | 1.38 [0.71; 2.05] |
| (6) Badde et al. (2014) - experiment 1 | 1.5312 | 0.4023 | 3.2% | 1.53 [0.74; 2.32] |
| (7) Badde et al. (2014) - experiment 2 | 2.0772 | 0.4913 | 2.4% | 2.08 [1.11; 3.04] |
| (8) Badde et al. (2015)- experiment 1 | 1.7541 | 0.5233 | 2.1% | 1.75 [0.73; 2.78] |
| (9) Badde et al. (2015)- experiment 2 | 1.9977 | 0.5437 | 2.0% | 2.00 [0.93; 3.06] |
| **Total (95% CI)** | | | **57.5%** | **1.50 [1.28; 1.72]** |
| Heterogeneity: Tau$^2$ = 0.0903; Chi$^2$ = 26.18, df = 17 (P = 0.0713); I$^2$ = 35.1% | | | | |
| **Eyes = closed** | | | | |
| (11) Cadieux & Shore (2013) - eyes closed, feet external | 1.6101 | 0.5583 | 1.9% | 1.61 [0.52; 2.70] |
| (12) Cadieux & Shore (2013) - eyes closed, hands | 1.3014 | 0.3333 | 4.2% | 1.30 [0.65; 1.95] |
| (16) Crollen, Albouy, et al. (2017) | 1.0885 | 0.4518 | 2.7% | 1.09 [0.20; 1.97] |
| (17) Crollen, Lazzouni, et al. (2017) | 0.9166 | 0.4217 | 3.0% | 0.92 [0.09; 1.74] |
| (18) Crollen et al. (2019) | 1.6086 | 0.4493 | 2.7% | 1.61 [0.73; 2.49] |
| (22) Kóbor et al. (2006) | 1.0878 | 0.3274 | 4.3% | 1.09 [0.45; 1.73] |
| (25) Nishikawa et al. (2015) | 1.7000 | 0.3873 | 3.4% | 1.70 [0.94; 2.46] |
| (26) Roberts & Humphreys (2008) | 1.1817 | 0.5462 | 2.0% | 1.18 [0.11; 2.25] |
| (27) Röder et al. (2004) - eyes closed | 1.6129 | 0.4957 | 2.3% | 1.61 [0.64; 2.58] |
| (29) Sambo et al. (2013) | 1.5964 | 0.5211 | 2.2% | 1.60 [0.57; 2.62] |
| (30) Schicke & Röder (2006) | 2.2107 | 0.7521 | 1.1% | 2.21 [0.74; 3.68] |
| (37) Unwalla, Cadieux, et al. (2021) - eyes closed | 1.3685 | 0.3419 | 4.1% | 1.37 [0.70; 2.04] |
| (39) Wada et al. (2004) | 2.0493 | 0.3323 | 4.2% | 2.05 [1.40; 2.70] |
| (4) Azañón, Mihaljevic, et al. (2016) | 0.9125 | 0.3296 | 4.3% | 0.91 [0.27; 1.56] |
| **Total (95% CI)** | | | **42.5%** | **1.39 [1.17; 1.61]** |
| Heterogeneity: Tau$^2$ = 0.0090; Chi$^2$ = 11.4, df = 13 (P = 0.5777); I$^2$ = 0% | | | | |
| **Total (95% CI)** | | | **100.0%** | **1.45 [1.30; 1.60]** |
| **Prediction interval** | | | | **[0.92; 1.97]** |
| Heterogeneity: Tau$^2$ = 0.0587; Chi$^2$ = 37.68, df = 31 (P = 0.1901); I$^2$ = 17.7% | | | | |
| Test for subgroup differences: Chi$^2$ = 0.54, df = 1 (P = 0.4626) | | | | |

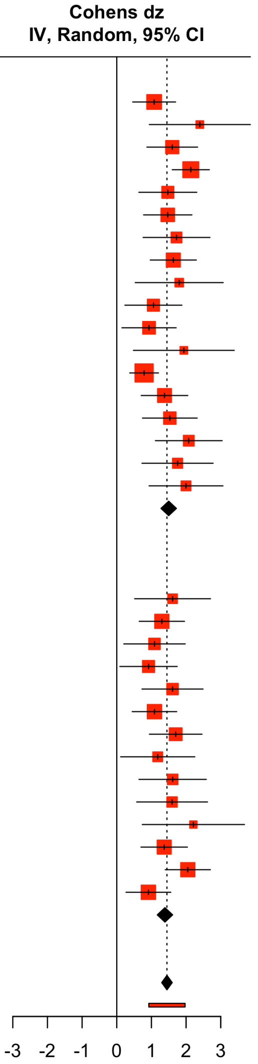


**Figure C2. Forest plot of the effect sizes of included studies grouped by the moderator visual input.** *SE* = standard error, CI = confidence interval, *df* = degrees of freedom. Red squares represent the point estimates for each study and black lines are 95% confidence intervals. The size of the squares reflects the study weight with which a study contributes to the pooled effect size. The black diamond symbolizes the pooled effect estimate, while the red line below denotes the prediction interval.

| Study or Subgroup | Cohens dz | SE | Weight | Cohens dz IV, Random, 95% CI |
|---|---|---|---|---|
| **Response.modality = hands** | | | | |
| (1) Azañón & Soto-Faraco (2007) - experiment 1 | 1.0784 | 0.3158 | 3.6% | 1.08 [0.46; 1.70] |
| (10) Badde et al. (2017) | 0.7224 | 0.3082 | 3.7% | 0.72 [0.12; 1.33] |
| (12) Cadieux & Shore (2013) - eyes closed, hands | 1.3014 | 0.3333 | 3.3% | 1.30 [0.65; 1.95] |
| (14) Cadieux & Shore (2013) - eyes open, hands | 1.6010 | 0.3733 | 2.9% | 1.60 [0.87; 2.33] |
| (16) Crollen, Albouy, et al. (2017) | 1.0885 | 0.4518 | 2.2% | 1.09 [0.20; 1.97] |
| (17) Crollen, Lazzouni, et al. (2017) | 0.9166 | 0.4217 | 2.4% | 0.92 [0.09; 1.74] |
| (19) Heed et al. (2012) - experiment 1 | 1.4733 | 0.4265 | 2.4% | 1.47 [0.64; 2.31] |
| (2) Azañón & Soto-Faraco (2007) - experiment 2 | 1.4715 | 0.3556 | 3.1% | 1.47 [0.77; 2.17] |
| (20) Heed et al. (2012) - experiment 2 | 1.7238 | 0.4916 | 1.9% | 1.72 [0.76; 2.69] |
| (21) Heed et al. (2016) | 1.6311 | 0.3384 | 3.3% | 1.63 [0.97; 2.29] |
| (25) Nishikawa et al. (2015) | 1.7000 | 0.3873 | 2.7% | 1.70 [0.94; 2.46] |
| (27) Röder et al. (2004) - eyes closed | 1.6129 | 0.4957 | 1.9% | 1.61 [0.64; 2.58] |
| (28) Röder et al. (2004) - eyes open | 1.0567 | 0.4198 | 2.5% | 1.06 [0.23; 1.88] |
| (29) Sambo et al. (2013) | 1.5964 | 0.5211 | 1.8% | 1.60 [0.57; 2.62] |
| (3) Azañón et al. (2015) | 0.9309 | 0.3995 | 2.6% | 0.93 [0.15; 1.71] |
| (30) Schicke & Röder (2006) | 2.2107 | 0.7521 | 1.0% | 2.21 [0.74; 3.68] |
| (32) Shore et al. (2002) | 1.9351 | 0.7422 | 1.0% | 1.94 [0.48; 3.39] |
| (33) Soto-Faraco & Azañón (2013) | 0.7888 | 0.2117 | 5.2% | 0.79 [0.37; 1.20] |
| (34) Unwalla et al. (2020) | 1.6435 | 0.3793 | 2.8% | 1.64 [0.90; 2.39] |
| (37) Unwalla, Cadieux, et al. (2021) - eyes closed | 1.3685 | 0.3419 | 3.2% | 1.37 [0.70; 2.04] |
| (38) Unwalla, Cadieux, et al. (2021) - eyes open | 1.3774 | 0.3431 | 3.2% | 1.38 [0.71; 2.05] |
| (39) Wada et al. (2004) | 2.0493 | 0.3323 | 3.3% | 2.05 [1.40; 2.70] |
| (4) Azañón, Mihaljevic, et al. (2016) | 0.9125 | 0.3296 | 3.4% | 0.91 [0.27; 1.56] |
| (5) Azañón, Radulova, et al. (2016) | 1.1880 | 0.4706 | 2.1% | 1.19 [0.27; 2.11] |
| (6) Badde et al. (2014) - experiment 1 | 1.5312 | 0.4023 | 2.6% | 1.53 [0.74; 2.32] |
| (7) Badde et al. (2014) - experiment 2 | 2.0772 | 0.4913 | 1.9% | 2.08 [1.11; 3.04] |
| (8) Badde et al. (2015)- experiment 1 | 1.7541 | 0.5233 | 1.8% | 1.75 [0.73; 2.78] |
| (9) Badde et al. (2015)- experiment 2 | 1.9977 | 0.5437 | 1.7% | 2.00 [0.93; 3.06] |
| **Total (95% CI)** | | | **73.5%** | **1.36 [1.20; 1.52]** |
| Heterogeneity: $Tau^2 = 0.0399$; $Chi^2 = 31.32$, df = 27 (P = 0.2581); $I^2 = 13.8\%$ | | | | |
| **Response.modality = feet external** | | | | |
| (11) Cadieux & Shore (2013) - eyes closed, feet external | 1.6101 | 0.5583 | 1.6% | 1.61 [0.52; 2.70] |
| (13) Cadieux & Shore (2013) - eyes open, feet external | 2.3970 | 0.7433 | 1.0% | 2.40 [0.94; 3.85] |
| (15) Cadieux et al. (2010) | 2.1362 | 0.2731 | 4.2% | 2.14 [1.60; 2.67] |
| (18) Crollen et al. (2019) | 1.6086 | 0.4493 | 2.2% | 1.61 [0.73; 2.49] |
| (23) Landry & Champoux (2018) | 2.4298 | 0.4982 | 1.9% | 2.43 [1.45; 3.41] |
| (26) Roberts & Humphreys (2008) | 1.1817 | 0.5462 | 1.6% | 1.18 [0.11; 2.25] |
| (31) Sharp et al. (2018) | 2.0924 | 0.5921 | 1.4% | 2.09 [0.93; 3.25] |
| **Total (95% CI)** | | | **14.0%** | **1.94 [1.55; 2.33]** |
| Heterogeneity: $Tau^2 = 0.0399$; $Chi^2 = 4.74$, df = 6 (P = 0.5780); $I^2 = 0\%$ | | | | |
| **Response.modality = feet anatomical** | | | | |
| (22) Kóbor et al. (2006) | 1.0878 | 0.3274 | 3.4% | 1.09 [0.45; 1.73] |
| (35) Unwalla, Goldreich, et al. (2021) - experiment 1 | 1.9387 | 0.4223 | 2.4% | 1.94 [1.11; 2.77] |
| (36) Unwalla, Goldreich, et al. (2021) - experiment 2 | 1.1193 | 0.2020 | 5.4% | 1.12 [0.72; 1.52] |
| **Total (95% CI)** | | | **11.3%** | **1.27 [0.27; 2.27]** |
| Heterogeneity: $Tau^2 = 0.0399$; $Chi^2 = 3.3$, df = 2 (P = 0.1916); $I^2 = 39.5\%$ | | | | |
| **Response.modality = verbal** | | | | |
| (24) Moseley et al. (2012) | 1.8025 | 0.6474 | 1.2% | 1.80 [0.53; 3.07] |
| **Total (95% CI)** | | | **100.0%** | **1.45 [1.30; 1.59]** |
| **Prediction interval** | | | | **[0.88; 2.01]** |
| Heterogeneity: $Tau^2 = 0.0715$; $Chi^2 = 52.38$, df = 38 (P = 0.0603); $I^2 = 27.5\%$ | | | | |
| Residual heterogeneity: $Tau^2 = 0.0399$; $Chi^2 = 39.37$, df = 35 (P = 0.2808); $I^2 = 11.1\%$ | | | | |
| Test for subgroup differences: $Chi^2 = 11.68$, df = 3 (P = 0.0086) | | | | |

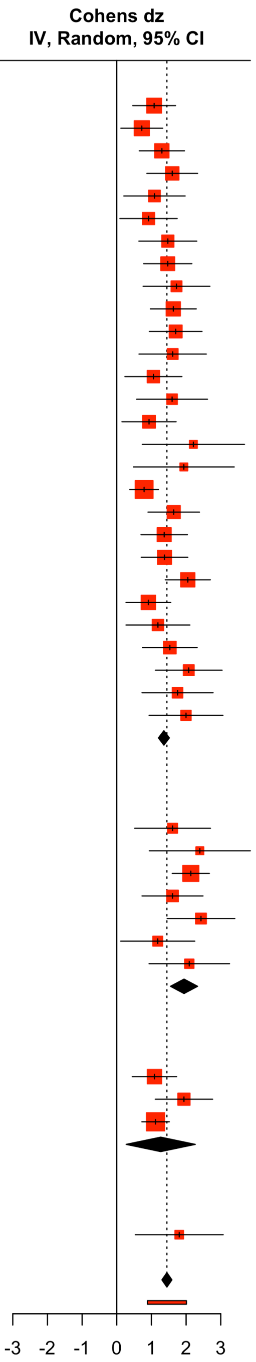


**Figure C3. Forest plot of the effect sizes of included studies grouped by the moderator response modality.** *SE* = standard error, CI = confidence interval, *df* = degrees of freedom. Red squares represent the point estimates for each study and black lines are 95% confidence intervals. The size of the squares reflects the study weight with which a study contributes to the pooled effect size. The black diamond symbolizes the pooled effect estimate, while the red line below denotes the prediction interval.

| Study or Subgroup | Cohens dz | SE | Weight | Cohens dz IV, Random, 95% CI |
|---|---|---|---|---|
| **Speed = unspeeded** | | | | |
| (1) Azañón & Soto-Faraco (2007) - experiment 1 | 1.0784 | 0.3158 | 4.0% | 1.08 [0.46; 1.70] |
| (19) Heed et al. (2012) - experiment 1 | 1.4733 | 0.4265 | 2.6% | 1.47 [0.64; 2.31] |
| (2) Azañón & Soto-Faraco (2007) - experiment 2 | 1.4715 | 0.3556 | 3.4% | 1.47 [0.77; 2.17] |
| (20) Heed et al. (2012) - experiment 2 | 1.7238 | 0.4916 | 2.1% | 1.72 [0.76; 2.69] |
| (22) Kóbor et al. (2006) | 1.0878 | 0.3274 | 3.8% | 1.09 [0.45; 1.73] |
| (23) Landry & Champoux (2018) | 2.4298 | 0.4982 | 2.0% | 2.43 [1.45; 3.41] |
| (26) Roberts & Humphreys (2008) | 1.1817 | 0.5462 | 1.7% | 1.18 [0.11; 2.25] |
| (27) Röder et al. (2004) - eyes closed | 1.6129 | 0.4957 | 2.1% | 1.61 [0.64; 2.58] |
| (28) Röder et al. (2004) - eyes open | 1.0567 | 0.4198 | 2.7% | 1.06 [0.23; 1.88] |
| (3) Azañón et al. (2015) | 0.9309 | 0.3995 | 2.9% | 0.93 [0.15; 1.71] |
| (30) Schicke & Röder (2006) | 2.2107 | 0.7521 | 1.0% | 2.21 [0.74; 3.68] |
| (31) Sharp et al. (2018) | 2.0924 | 0.5921 | 1.5% | 2.09 [0.93; 3.25] |
| (32) Shore et al. (2002) | 1.9351 | 0.7422 | 1.0% | 1.94 [0.48; 3.39] |
| (4) Azañón, Mihaljevic, et al. (2016) | 0.9125 | 0.3296 | 3.8% | 0.91 [0.27; 1.56] |
| (5) Azañón, Radulova, et al. (2016) | 1.1880 | 0.4706 | 2.2% | 1.19 [0.27; 2.11] |
| (8) Badde et al. (2015)- experiment 1 | 1.7541 | 0.5233 | 1.9% | 1.75 [0.73; 2.78] |
| (9) Badde et al. (2015)- experiment 2 | 1.9977 | 0.5437 | 1.8% | 2.00 [0.93; 3.06] |
| **Total (95% CI)** | | | **40.7%** | **1.38 [1.15; 1.60]** |
| Heterogeneity: $Tau^2$ = 0.0021; $Chi^2$ = 16.17, df = 16 (P = 0.4414); $I^2$ = 1% | | | | |
| **Speed = time restriction** | | | | |
| (10) Badde et al. (2017) | 0.7224 | 0.3082 | 4.2% | 0.72 [0.12; 1.33] |
| (11) Cadieux & Shore (2013) - eyes closed, feet external | 1.6101 | 0.5583 | 1.7% | 1.61 [0.52; 2.70] |
| (12) Cadieux & Shore (2013) - eyes closed, hands | 1.3014 | 0.3333 | 3.8% | 1.30 [0.65; 1.95] |
| (13) Cadieux & Shore (2013) - eyes open, feet external | 2.3970 | 0.7433 | 1.0% | 2.40 [0.94; 3.85] |
| (14) Cadieux & Shore (2013) - eyes open, hands | 1.6010 | 0.3733 | 3.2% | 1.60 [0.87; 2.33] |
| (15) Cadieux et al. (2010) | 2.1362 | 0.2731 | 4.8% | 2.14 [1.60; 2.67] |
| (16) Crollen, Albouy, et al. (2017) | 1.0885 | 0.4518 | 2.4% | 1.09 [0.20; 1.97] |
| (17) Crollen, Lazzouni, et al. (2017) | 0.9166 | 0.4217 | 2.7% | 0.92 [0.09; 1.74] |
| (18) Crollen et al. (2019) | 1.6086 | 0.4493 | 2.4% | 1.61 [0.73; 2.49] |
| (25) Nishikawa et al. (2015) | 1.7000 | 0.3873 | 3.0% | 1.70 [0.94; 2.46] |
| (29) Sambo et al. (2013) | 1.5964 | 0.5211 | 1.9% | 1.60 [0.57; 2.62] |
| (34) Unwalla et al. (2020) | 1.6435 | 0.3793 | 3.1% | 1.64 [0.90; 2.39] |
| (35) Unwalla, Goldreich, et al. (2021) - experiment 1 | 1.9387 | 0.4223 | 2.7% | 1.94 [1.11; 2.77] |
| (36) Unwalla, Goldreich, et al. (2021) - experiment 2 | 1.1193 | 0.2020 | 6.6% | 1.12 [0.72; 1.52] |
| (37) Unwalla, Cadieux, et al. (2021) - eyes closed | 1.3685 | 0.3419 | 3.6% | 1.37 [0.70; 2.04] |
| (38) Unwalla, Cadieux, et al. (2021) - eyes open | 1.3774 | 0.3431 | 3.6% | 1.38 [0.71; 2.05] |
| (39) Wada et al. (2004) | 2.0493 | 0.3323 | 3.8% | 2.05 [1.40; 2.70] |
| (6) Badde et al. (2014) - experiment 1 | 1.5312 | 0.4023 | 2.9% | 1.53 [0.74; 2.32] |
| (7) Badde et al. (2014) - experiment 2 | 2.0772 | 0.4913 | 2.1% | 2.08 [1.11; 3.04] |
| **Total (95% CI)** | | | **59.3%** | **1.51 [1.31; 1.72]** |
| Heterogeneity: $Tau^2$ = 0.0715; $Chi^2$ = 25.95, df = 18 (P = 0.1009); $I^2$ = 30.6% | | | | |
| **Total (95% CI)** | | | **100.0%** | **1.47 [1.32; 1.62]** |
| **Prediction interval** | | | | **[0.98; 1.96]** |
| Heterogeneity: $Tau^2$ = 0.0526; $Chi^2$ = 42.77, df = 35 (P = 0.1721); $I^2$ = 18.2% | | | | |
| Test for subgroup differences: $Chi^2$ = 0.89, df = 1 (P = 0.3463) | | | | |

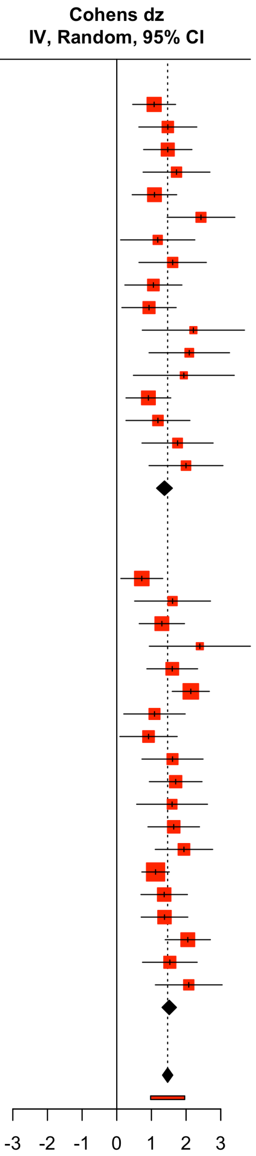


**Figure C4. Forest plot of the effect sizes of included studies grouped by the moderator task instructions of speed.** *SE* = standard error, CI = confidence interval, *df* = degrees of freedom. Red squares represent the point estimates for each study and black lines are 95% confidence intervals. The size of the squares reflects the study weight with which a study contributes to the pooled effect size. The black diamond symbolizes the pooled effect estimate, while the red line below denotes the prediction interval.
